# NFIA facilitates CHD4 recruitment to activate fatty acid metabolic genes in postnatal cardiomyocytes

**DOI:** 10.64898/2026.11.23.767975

**Authors:** Mingqi Cai, Yan Y Sweat, Yu-Wen Chen, Liu Mei, Koutakis, Mason E Sweat, Wei Shi

**Affiliations:** Mary & Dick Holland Regenerative Medicine Center, University of Nebraska Medical Center, Omaha, NE 68198, USA; Division of Cardiovascular Medicine, Department of Internal Medicine, University of Nebraska Medical Center, Omaha, NE 68198, USA; Moss Heart Center, Division of Cardiology, Department of Internal Medicine. UT Southwestern Medical Center, Dallas, TX 75390, USA; Department of Biochemistry and Molecular Biology, University of Nebraska Medical Center, Omaha, NE 68198, USA; Department of Cellular and Integrative Physiology, University of Nebraska Medical Center, Omaha, NE 68198, USA

**Keywords:** CHD4, NFIA, postnatal cardiomyocytes, fatty acid metabolism, gene activation

## Abstract

**BACKGROUND:** Mammalian cardiomyocyte maturation involves a postnatal metabolic shift from glycolysis toward fatty acid oxidation, yet the transcriptional mechanisms underlying this transition remain incompletely understood. Chromodomain Helicase DNA-binding protein 4 (CHD4), the catalytic core of the Nucleosome Remodeling and Deacetylase (NuRD) complex, is essential for postnatal cardiac homeostasis and mitochondrial function. However, whether CHD4 regulates the transcriptional programs underlying postnatal cardiomyocyte metabolic maturation remains unknown.

**METHODS:** Adeno-associated virus (AAV)-mediated *Cre* recombination was used to generate postnatal cardiomyocyte-specific *Chd4-*deficient mice. Cardiac function, mitochondrial ultrastructure and function, and lipid accumulation were assessed. Integrated analyses of embryonic and postnatal cardiac transcriptomes, CHD4 chromatin occupancy, and immunoprecipitation-coupled proteomics were performed to identify CHD4-regulated metabolic genes and interacting transcription factors. CUT&RUN (Cleavage Under Targets and Release Using Nuclease) assays in primary cardiomyocytes were used to investigate the role of Nuclear Factor I A (NFIA) in recruiting CHD4 to fatty acid metabolic gene loci.

**RESULTS:** Postnatal cardiomyocyte-specific *Chd4* deletion resulted in cardiac dysfunction, altered lipid distribution, disrupted mitochondrial cristae architecture, premature mortality, and downregulation of fatty acid metabolic genes. Integrated transcriptomic and chromatin occupancy analyses revealed that CHD4 activates a gene program in postnatal cardiomyocytes enriched in fatty acid metabolic pathways. Proteomic profiling identified NFIA as a postnatal-specific CHD4-interacting transcription factor. A subset of postnatal CHD4-bound genes that were downregulated following *Chd4* deletion also were reduced in cardiomyocyte-specific *Nfia-*deficient hearts. Furthermore, *Nfia* depletion reduced CHD4 occupancy at postnatal fatty acid metabolic gene loci, supporting an NFIA-dependent CHD4 recruitment mechanism in postnatal cardiomyocytes.

**CONCLUSIONS:** These findings identify NFIA as a developmental regulator that facilitates CHD4 recruitment to genomic regions in postnatal cardiomyocytes, contributing to the activation of fatty acid metabolic genes during cardiac maturation.

**Novelty and Significance:** *What Is Known?:*

- Postnatal cardiomyocyte maturation involves a metabolic transition from glycolysis toward fatty acid oxidation to meet the increasing energetic demands of the heart.
- CHD4, the catalytic core of the NuRD complex, regulates embryonic cardiac development and is predominantly associated with transcriptional repression.
- Transcription factor NFIA regulates fatty acid metabolic gene expression in noncardiac tissues, but its role in cardiac metabolic maturation remains unknown.

*What New Information Does This Article Contribute?:*

- CHD4 activates a gene program in postnatal cardiomyocytes promoting fatty acid metabolism, and postnatal cardiomyocyte-specific CHD4 deficiency disrupts myocardial lipid handling.
- NFIA physically associates with CHD4 in postnatal cardiomyocytes, and NFIA deficiency downregulates a subset of CHD4-bound genes that are also downregulated following *Chd4* deletion.
- *Nfia* depletion reduces CHD4 occupancy at selected postnatal-specific regulatory regions of fatty acid metabolic genes, supporting an NFIA-dependent mechanism of CHD4 recruitment. Our findings identify a developmentally regulated NFIA-CHD4 mechanism that promotes the expression of fatty acid metabolic genes during postnatal cardiomyocyte maturation. This work reveals an uncharacterized gene-activating function of CHD4 in the postnatal heart and expands our understanding of the transcriptional mechanisms underlying cardiac metabolic maturation.

## INTRODUCTION

During the early postnatal transition, mammalian cardiomyocytes undergo cellular maturation processes characterized by physiological hypertrophy, loss of regenerative capacity, and metabolic pathway switch^1,2^. In the embryonic heart, energy production relies primarily on glycolysis to sustain cardiac growth in a relatively hypoxic environment. Following birth, increased oxygen availability and the transition to lipid-rich nutrition promote a shift toward fatty acid oxidation and oxidative phosphorylation, supporting the increased energetic demands of the postnatal heart^1–8^. While the physiological drivers of this metabolic transition are intensively studied, transcriptional networks regulating postnatal cardiomyocyte metabolic maturation remain poorly understood.

Chromodomain Helicase DNA-binding protein 4 (CHD4), a core catalytic ATPase subunit of the Nucleosome Remodeling and Deacetylase (NuRD) complex^9–14^, is an essential chromatin remodeler that regulates cardiac gene transcription and maintains cardiomyocyte identity and homeostasis^15–22^. Pathogenic variants in CHD4 have been associated with congenital heart defects, including atrial and ventricular septal defects and Tetralogy of Fallot^23–26^. CHD4 has been traditionally recognized as a transcriptional repressor through its association with histone deacetylases (HDAC1/2)^9,13,14^. However, emerging evidence suggests that CHD4 acts as both a transcriptional repressor and an activator, depending on its partner transcription factors and local epigenetic environment^27^. Consistent with this mechanism, we recently identified that in postnatal atrial cardiomyocytes, T-box transcription factor 5 (TBX5) recruits CHD4 to specific enhancers to maintain accessibility; conversely, inactivating CHD4 results in the loss of TBX5 occupancy and genomic accessibility, leading to enhancer inactivation and gene downregulation^21^. These findings suggest that CHD4 functions as a context-dependent regulator of transcriptional states in the heart. Previous studies established that CHD4 directly targets promoter regions to activate selected mitochondrial genes in striated muscle (e.g., *Mipep, Pgc1a, Sirt5*, and *Ucp2*)^17^. However, whether CHD4 drives the broader postnatal activation of fatty acid metabolic networks in cardiomyocytes, and whether its gene-activating function is stage-specific, have not been determined. Moreover, CHD4 relies on transcription factors to guide its binding to specific genomic loci as it lacks DNA-binding motifs. For example, we previously identified that CHD4 is recruited by TBX5, GATA-Binding Factor 4 (GATA4), and NK2 Homeobox 5 (NKX2-5) to access and inactivate non-cardiac myofibril gene loci in embryonic cardiomyocytes^18^. However, how CHD4 is recruited to fatty acid metabolic gene loci in postnatal cardiomyocytes remain poorly studied.

Nuclear Factor I A (NFIA), a member of the Nuclear Factor I family, has been to cooperates with peroxisome proliferator-activated receptor gamma (PPARγ) at brown-fat-specific enhancers, facilitating PPARγ occupancy and promoting transcriptional programs associated with brown adipocyte differentiation and mitochondrial metabolism^28^. Despite its established roles in regulating lipid metabolism in adipocytes and other organ systems (e.g., articular chondrocytes and spinal motor neuron)^28–33^, the function of NFIA in the postnatal mammalian heart has not been investigated.

In this study, using cardiomyocyte-specific knockout mouse models and integrated multi-omic analyses, we uncovered a novel mechanism by which NFIA facilitates CHD4 recruitment to postnatal-specific regulatory loci, thereby activating the fatty acid metabolic gene program during postnatal cardiac maturation.

## MATERIALS AND METHODS

### Mice

The *Chd4^flox^* allele was kindly provided by Dr. Frank Conlon’s laboratory (originally generated by Dr. Katia Georgeopolos’s laboratory^34^). The *Nfia^flox^* allele was obtained from Dr. Benjamin Deneen’s laboratory^35^. The Tg(Tnnt2-cre)5Blh/JiaoJ (RRID: IMSR_JAX:024240) allele^36^ and C57Bl/6 mice were purchased from The Jackson Laboratory. CD-1 mice were purchased from Charles River Laboratories. All transgenic mouse strains were maintained on a mixed genetic background. Genotyping was performed as previously described^22,37,38^ or per literature^35^. Mice were housed under standard 12-hour light/dark cycles with free access to standard lab chow and water. The pAAV9-cTNT-Cre-eGFP and pAAV9-cTNT-eGFP viral particles were subcutaneously injected to neonatal *Chd4^flox/flox^* pups at postnatal day 1 (P1) at 2×10^11^ viral genome/pup (30-50 μl/pup depending on the viral particle titration). At 4 weeks post viral treatment, the *Chd4-cmKO* (pAAV9-cTNT2-Cre-eGFP::*Chd4^flox/flox^*) and *Control* (pAAV9-cTNT2-eGFP::*Chd4^flox/flox^*) mice were subjected to the record of voluntary wheel-running activity with the Scurry Activity Wheel Chamber System, echocardiographic measurement, and sampling. The Institutional Animal Care and Use Committee (IACUC; Protocol# 24-042-07) at the University of Nebraska Medical Center (UNMC) approved all animal studies.

### Voluntary wheel-running activity

Voluntary physical activity was assessed in 4-week-old *Chd4-cmKO* and *Control* mice using the Scurry Activity Wheel Chamber System (Lafayette Instrument) at the UNMC Animal Behavior Core Facility. Mice were individually housed in activity wheel chambers maintained on a 12-hour light/dark cycle with ad libitum access to water and powdered standard chow. Food intake was measured by weighing the powdered chow before and after the testing period. Wheel revolutions were continuously recorded over 24 hours using a lid-mounted Scurry Mouse Activity Counter and Monitor and quantified using Scurry Activity Monitoring Software.

### Echocardiography

Transthoracic two-dimensional (2D) echocardiography was performed using the VisualSonics Vevo 3100 ultrasound system equipped with the MX550S transducer (FUJIFILM VisualSonics, Toronto, ON, Canada). Mice were anesthetized in an induction chamber with 2% isoflurane and then placed supine on a heated ECG board, with limbs secured to ECG pads coated with electrode gel. Anesthesia was maintained via mask delivering 2% isoflurane. Applied ultrasound gel to the mouse chest and then adjusted the platform and probe to appropriate angles and positions to show long-axis and short-axis views of the left ventricle (LV) using B-mode. And M-mode was then applied to capture dynamic long-axis and short-axis recordings across the cardiac cycle and the parameters including heart rate, ejection fraction, fractional shortening, stroke volume, cardiac output, LV end-diastolic internal diameter (Diameter;d), and LV mass were derived from primary measurements using Vevo LAB software. Pulmonary artery velocity time integral (PA-VTI) and aortic valve velocity time integral (AVo-VTI) were assessed using Pulse Wave Doppler echocardiography. Doppler flow velocity waveforms were recorded, and the systolic velocity profiles were traced to calculate the velocity time integral (VTI) for each measurement. Pulsed-wave Doppler echocardiography was also used to assess left and right ventricular diastolic function by measuring peak early diastolic transmitral or transtricuspid inflow velocity (E wave) during passive ventricular filling and peak late diastolic inflow velocity (A wave) during atrial contraction. The E/A ratio represents one important parameter for analyzing diastolic function. All echo procedures and data analyses were performed in a blinded manner, with operators and analysts unaware of the treatment or groupings of each mouse.

### RNA extraction

Total RNA was extracted from heart tissue as previously described^21^ with minor modifications. Briefly, hearts were perfused with 15 mL of ice-cold PBS, dissected, minced into small pieces, and stored in RNAlater Stabilization Solution (Thermo Fisher Scientific, Cat# AM7020) until processing. Tissue samples were homogenized in 1 mL of ice-cold TRIzol Reagent (Thermo Fisher Scientific, Cat# 15596026) and mixed with 0.2 mL of chloroform for phase separation. Following centrifugation, the aqueous phase was transferred, and total RNA was purified using the PureLink RNA Mini Kit (Thermo Fisher Scientific, Cat#12183018A) according to the manufacturer’s instructions. DNA was removed by incubating the eluate with PureLink DNase (Thermo Fisher Scientific, Cat#12185010) at room temperature for 15 min. RNA concentration and purity were measured using a NanoDrop spectrophotometer (Thermo Fisher Scientific), and RNA integrity was evaluated using the Qubit RNA IQ Assay Kit (Thermo Fisher Scientific, Cat#Q33221). Samples with RNA integrity numbers between 8.0 and 10.0 were used for bulk RNA-sequencing.

### RNA-sequencing and bioinformatic analysis

Library prep and sequencing were performed at AZENTA Life Sciences (Burlington, MA, US) as previously described^21^. Briefly, poly-A selection was performed to enrich mRNA followed by mRNA fragmentation, random priming, and cDNA synthesis. The double-stranded DNA then underwent end repair, 5’ phosphorylation, dA-tailing, adaptor ligation, and barcode-PCR. Libraries were sequenced (2×150 Paired End (PE); Illumina HiSeq 2500) to a target depth of >30 million reads. Sequence reads were trimmed to remove possible adapter sequences and nucleotides with poor quality using Trimmomatic v.0.36. The trimmed reads were mapped to the Mus musculus GRCm38 reference genome available on ENSEMBL using the STAR aligner v.2.5.2b. Unique gene hit counts were calculated by using featureCounts from the Subread package v.1.5.2. The hit counts were summarized and reported using the gene_id feature in the annotation file. Only unique reads that fell within exon regions were counted. After extraction of gene hit counts, the gene hit counts table was used for downstream differential expression analysis with DESeq2. The Wald test was used to generate p-values and log2 fold changes. Genes with an adjusted p-value < 0.05 and absolute log2 fold change > 0.5 were called as differentially expressed genes (DEGs) for comparison. Gene Ontology (GO) analysis was performed using the Panther.db. Classification System (PANTHER 19.0)^39^ and the Gene Set Enrichment Analysis software as previously described^19^.

### RT-qPCR

Reverse transcription was performed by using Applied Biosystem ^TM^ High-Capacity cDNA Reverse Transcription Kit (Thermo Fisher Scientific, Cat# 4374966) following the manufacturer’s instructions to make cDNA. Quantitative PCR (qPCR) was performed using PowerUP SYBR Green master mix (Thermo Fisher Scientific, Cat# 25742) in a QuantStudio^TM^ 3 real-time PCR system (Thermo Fisher Scientific, Applied Biosystems) at the following thermocycling conditions:

|  |  |  |  |
|---|---|---|---|
| Hold Stage | Step 1 | 50 °C | 2 min |
|  | Step 2 | 95 °C | 2 min |
| PCR Stage (40 cycles) | Step 1 | 95 °C | 15 s |
|  | Step 2 | 60 °C | 15 s |
|  | Step 3 | 72 °C | 1 min |
| Melt Curve Stage | Step 1 | 95 °C | 15 s |
|  | Step 2 | 60 °C | 1 min |
|  | Step 3 | 95 °C | 15 s |

The mRNA level was calculated by the relative quantitative method 2^(−δδCt) with the formula: δδCt = δCt(Experimental group) − δCt(control group), δCt = Ct(target gene) − Ct(internal reference), and 2^(−δδCt) represented the mRNA relative transcriptional level. Primers for amplifying are listed below:

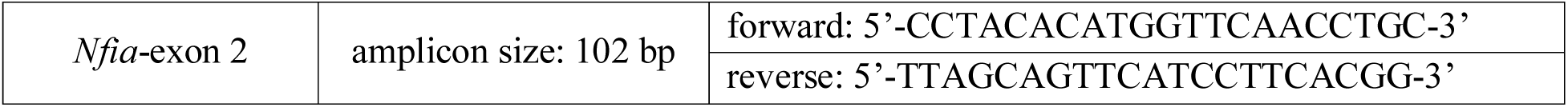

### Primary cardiomyocyte isolation

Isolation of embryonic and postnatal mouse cardiomyocytes was performed as previously reported^20^. Briefly, time-mated female wild type CD1 mice were dissected at E11.5 and embryonic hearts were isolated and rinsed with pre-warmed PBS and culture media sequentially. Wild type CD1 or *Nfia^fl/fl^* neonatal mice were euthanized on postnatal day 0 (P0) and the hearts were harvested. Embryonic or postnatal hearts were placed in 2-mL tubes containing 500 µL of 0.25% trypsin-EDTA (Thermo Fisher Scientific, Cat# 25200056) and digested with rotation at 37 °C for 5 min. 1 mL pipette with filter tips were used to pipette the tissue up and down several time after every 5 min of incubation. The solution from first round of digestion was discarded because most cells were erythrocytes. 500 μL fresh trypsin-EDTA (0.25%) was then added into each tube for the following rounds of digestion. After 5 min, the solution (excluding the heart tissues) was collected in a new 15 mL tube with culture media containing 10% FBS in DMEM to inactivate trypsin. The process was repeated about six times until all the tissue was digested. All solutions that contain dissociated cells were combined and filtered through 100 µm cell strainers and spun at 1000 x g for 10 min at room temperature. The supernatant solution was aspirated, and the cell pellet was resuspended with culture media. The resuspended cells were seeded in a 6-cm dish for 1 hour to allow those non-cardiomyocytes to attach to the plate while most cardiomyocytes remained suspended in the culture media. The unattached cells were then transferred to the dishes precoated with 0.2% gelatin, and the dishes were gently shaken to ensure even cell distribution.

### Cleavage Under Targets & Release Using Nuclease (CUT&RUN) and Library Preparation

CUT&RUN was performed in E11.5 and P0 isolated cardiomyocytes following the user manual of CUTANA^TM^CHIC / CUT&RUN kit (EpiCypher, Cat#14-1048). Briefly, ConA beads were activated, followed by binding of the isolated cardiomyocytes to the activated beads and subsequent cell permeabilization. Primary antibodies (CHD4, 0.5 µg, EpiCypher, Cat# 13-2016; NFIA, 0.5 µg, Sigma-Aldrich, Cat# HPA006111; IgG, 0.5 µg, EpiCypher, Cat# 13-0042) were added to incubate the cells overnight at 4 °C on a nutator. The next day, pAG-MNase was added to each reaction and then digested and released targeted chromatin. After DNA purification, DNA was quantified using the Qubit fluorometer and 1X dsDNA HS Assay Kit (Thermo Fisher Scientific, Cat# Q33230). DNA samples were then ready for library preparation following the instructions of CUT&RUN Library Prep Kit (EpiCypher, Cat# 14-1001-48). Briefly, 5 ng CUT&RUN-enriched DNA samples went through end repair, adaptor ligation and U-excision, DNA cleanup, indexing PCR with unique pair of primers, and PCR cleanup. 1 µL purified CUT&RUN library was quantified using the Qubit fluorometer with the 1X dsDNA HS Assay Kit per manufacturer’s instructions (Thermo Fisher Scientific, Cat# Q33230).

### CUT&RUN-qPCR

CUT&RUN libraries were normalized to the same concentration and were diluted 1:1 with nuclease-free H2O. Genomic DNA from neonatal cardiomyocytes was purified as the input control and was diluted in 1:100. The diluted CUT&RUN libraries and input DNA were then used for qPCR with the same thermocycling conditions described above with the following primers:

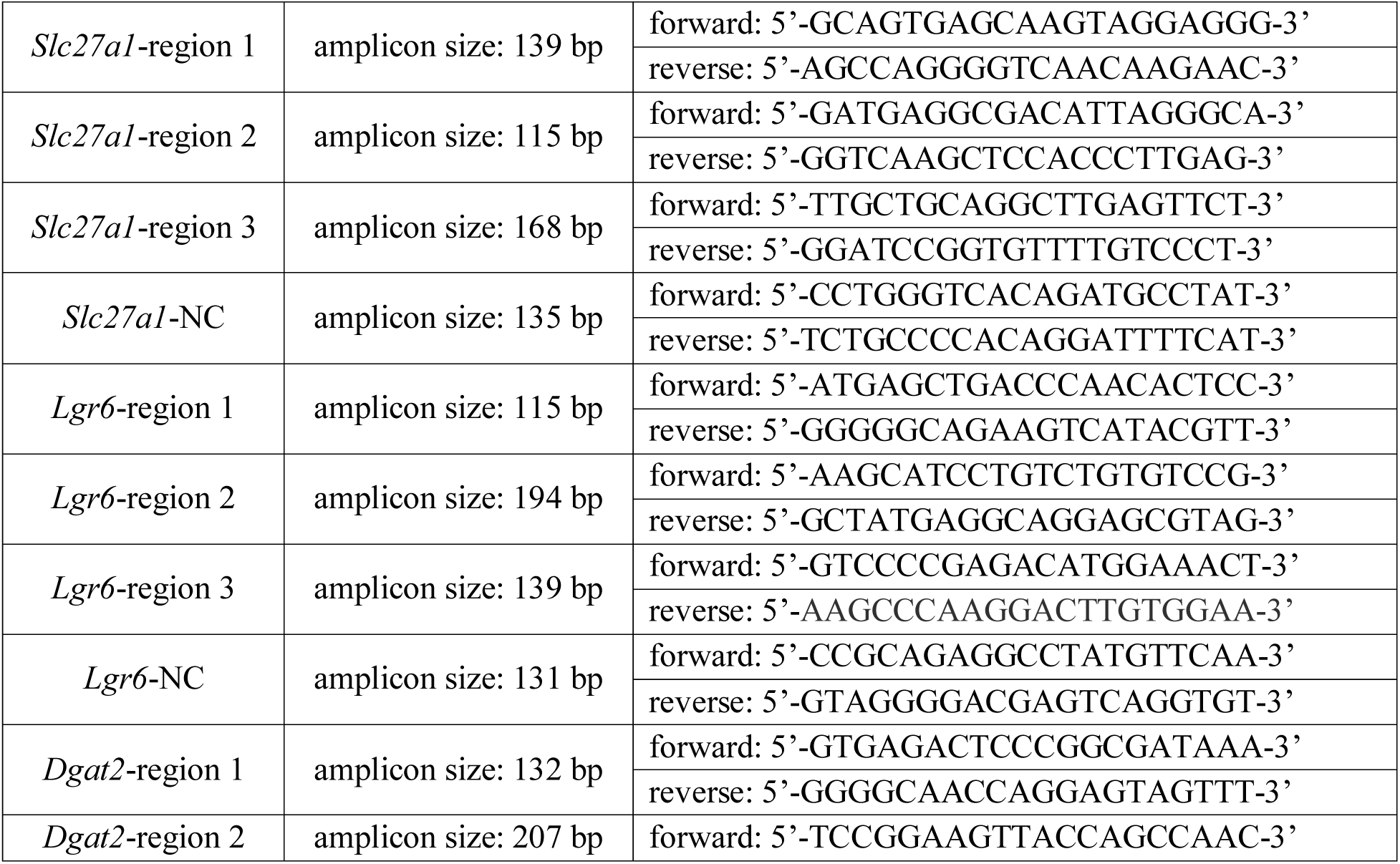
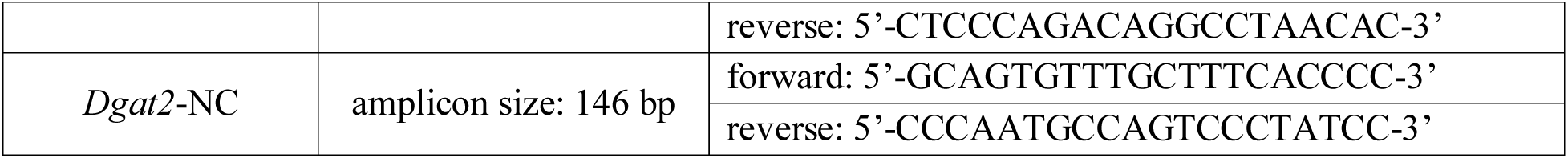

#### Enrichment calculations

Original Ct values (Ori-Ct) were corrected by dilution factors.

Corrected Ct values for CUT&RUN samples (Corr-Ct(C&R)) = Ori-Ct(C&R)-Log_2_2

Corrected Ct values for input DNA (Corr-Ct(input)) = Ori-Ct(input)-Log_2_100

δCt = [Corr-Ct(C&R)] – [Corr-Ct(input)]

The enrichment (%) over genomic DNA background (input) at each region = 2^ (-δCt)×100%

Then we calculated the enrichment fold change over each gene locus’s selected negative control (NC) region: Target Region Enrichment (over input) / NC Enrichment (over input) = Fold Change over NC, and the values of which were used for plotting and statistical analyses.

### Immunohistochemistry

Immunohistochemistry staining and imaging were performed as previously reported^19^. For heart tissue, paraffin-embedded sections were deparaffinized and rehydrated, and cryosections were heated at 55 °C for 20 min before staining. Sections were then immersed in boiled antigen retrieval buffer (10 mM Sodium Citrate, 0.05% Tween 20, pH 6.0) for 20 min, the cooled down sections were blocked with blocking buffer (10% normal goat serum (Abcam, Cat# ab7481), 1% Triton X-100 in PBS) at room temperature for 1 hour, and then incubated with desired primary antibody solution diluted in 10% blocking buffer (SLC27A1, 1:100, Thermo Fisher Scientific, Cat# PA5-110387; Cd36, 1:200, Thermo Fisher Scientific, Cat# PA1-16813; CHD4, 1:100, Active Motif, Cat# 39695; Tropomyosin, 1:50, Developmental Studies Hybridoma Bank, Cat# CH1; Troponin T, 1:200, Abcam, Cat# ab209813; NFIA, 1:250, Sigma-Aldrich, HPA006111) in a humidified chamber at 4 °C overnight. The next day, the sections were washed three times with washing buffer (10% blocking buffer) and then incubated with fluorescence-conjugated secondary antibodies (Goat anti-Rabbit IgG (H+L) Highly Cross-Adsorbed Secondary Antibody, Alexa Fluor™ 546, Thermo Fisher Scientific, Cat# A-11035; Goat anti-Mouse IgG (H+L) Cross-Adsorbed Secondary Antibody, Alexa Fluor™ 488, Thermo Fisher Scientific, Cat# A-11001) that were diluted in washing buffer (1:1000) at room temperature for 1 hour. After three times of washing, sections were counterstained with DAPI (Thermo Fisher Scientific, Cat# 62248) and mounted in Fluoromount-G^TM^ Mounting Medium (Thermo Fisher Scientific, Cat# 00-4958-02). Images were captured on a Zeiss LSM 880 laser scanning confocal microscope.

### Oil Red O staining

Oil Red O solution (Sigma-Aldrich, Cat# O1391) was diluted with distilled water to prepare a 60% (v/v) working solution, allowed to stand for 10 min, and filtered through 0.22 μM filter before use. Cryosections were immersed in 60% isopropanol for 2 min and excess solution was removed. The sections were then incubated with Oil Red O working solution for 20 min at room temperature, followed by immersion in 60% isopropanol for 1 min and rinsing with distilled water. Then the slides were mounted using Fluoromount-G^TM^ Mounting Medium (Thermo Fisher Scientific, Cat# 00-4958-02), covered with coverslips, and ready for brightfield scanning.

### High-resolution respirometry of fresh cardiac tissue (Oroboros)

Mitochondrial respiration in freshly isolated cardiac tissue was assessed by high-resolution respirometry using an Oroboros O2k system at 37°C. Fresh cardiac tissue was mechanically permeabilized and maintained under hyperoxygenated conditions (300–450 μM O₂) to ensure adequate oxygen diffusion throughout the tissue during measurements. Respiration was measured using a substrate-uncoupler-inhibitor titration (SUIT) protocol based on SUIT-002, with minor modifications. Briefly, 7.5 mM adenosine diphosphate (ADP; Sigma, 5285) was added to stimulate oxidative phosphorylation (OXPHOS) supported by endogenous substrates. Subsequently, 0.3 mM malate (Sigma, M1000) and 0.5 mM octanoylcarnitine (Tocris, 0605) were added to assess OXPHOS capacity supported by fatty acid oxidation (FAO). Cytochrome c (10 μM; Sigma, C7752) was then added to evaluate mitochondrial outer membrane integrity. Samples exhibiting an increase in oxygen consumption of 20% or greater following cytochrome c addition were excluded from the analysis. Additional substrates, including malate (2 mM), pyruvate (5 mM; Sigma, P2256), and glutamate (10 mM; Sigma, G1626), were subsequently added to assess combined FAO-and complex I (CI)-supported OXPHOS respiration. Succinate (10 mM; Sigma, S2378) was then added to evaluate respiration supported by FAO, CI, and complex II (CII). Next, carbonyl cyanide m-chlorophenyl hydrazone (CCCP; Sigma, C2759) was titrated stepwise (1 μL of 0.1 μM per injection) until maximal stable oxygen consumption was achieved, allowing assessment of maximal electron transfer system (ETS) capacity under uncoupled conditions. Finally, rotenone (1 μM; Sigma, R8875) and antimycin A (5 μM; Sigma, A8674) were sequentially added to inhibit mitochondrial electron transport and determine residual oxygen consumption (ROX), representing non-mitochondrial oxygen consumption. Mitochondrial oxygen consumption rates were calculated by subtracting ROX from the measured oxygen flux, normalized to wet tissue weight, and expressed as pmol O₂·s⁻¹·mg⁻¹.

### Cardiac nuclei extraction

Frozen hearts from P1-P2 wildtype C57BL/6 mouse hearts were homogenized using a mortar and pestle in liquid nitrogen. Nuclei were extracted as previously described^40^, and snap-frozen in liquid nitrogen and then stored it in-80 °C freezer before immunoprecipitation (for Figure 5B). For NFIA-IP/MS, the cardiac nuclei were isolated and were magnetic-activated cell sorting (MACS) for PCM1^+^ using protocols from Quaife-Ryan et al.^41^ and Wang et al.^42^, respectively. Nuclei collected before and after sorting were stained and examined by fluorescence microscopy to evaluate the enrichment of PCM1⁺ nuclei (Figure S5B). The PCM1⁺ nuclei were used for NFIA-IP/MS.

### Immunoprecipitation

Isolated cardiac nuclei samples from P1-P2 wildtype C57BL/6 mouse hearts were conjugated with rabbit anti-CHD4 (Abcam, Cat# ab72418), or rabbit anti-NFIA (Sigma-Aldrich, Cat# HPA006111), or negative control rabbit anti-IgG (Sigma-Aldrich, Cat# I5006) antibodies to magnetic beads (Thermo Fisher Scientific, Cat#10003D and #10001D) as previously described^19^. 5% lysate without adding antibodies was used as input controls. The immuno-isolated proteins were then subjected to Western Blot analysis or Mass Spectrometry analysis.

### Mass spectrometry

Immunoprecipitation (IP) elutions were solubilized in S-Trap lysis buffer containing SDS and Benzonase. Proteins were reduced with tris(2-carboxyethyl)phosphine (TCEP), alkylated with acrylamide, and transferred onto an S-Trap Turbo column for automated contaminant removal and enzymatic digestion with a Trypsin/Lys-C protease mix. Following digestion, peptides were eluted, collected, and quantified using the Pierce Fluorometric Peptide Assay. Internal retention time standards were spiked into each sample, and peptides were injected directly without normalization to total peptide amount. Samples were analyzed by liquid chromatography–tandem mass spectrometry (LC-MS/MS) on an Orbitrap Astral mass spectrometer operating in data-independent acquisition (DIA) mode. Raw MS data were processed and searched against the Mus musculus (mouse) reference proteome using Spectronaut, specifying methionine oxidation and cysteine propionamide as variable and fixed modifications, respectively.

To reduce the chances of false discovery and bolster protein level identification confidence, the raw results were filtered to require at least 2 peptide identifications per protein. All reported identifications were assigned to high FDR confidence by the search algorithm. Raw data was sequentially processed with Spectronaut and MsStats to be normalized to reduce technical or systematic variations and to improve statistical power.

### Western blot analyses

For extracting total myocardial proteins, frozen heart tissue was homogenized using a mortar and pestle. Heart homogenate was then lysed on ice for 30 min using RIPA lysis buffer (150 mM Sodium Chloride, 50 mM Tris-HCl (pH 7.4), 1% NP-40, 0.5% Sodium deoxycholate, and 0.1% SDS) with 1x Halt™ Protease and Phosphatase Inhibitor Cocktail (Thermo Fisher Scientific, Cat# 78446). The samples were then spun at 12000 rpm at 4 °C for 15 min. The supernatant was collected as total protein extracts from heart tissue. The protein concentration was quantified by Bicinchoninic acid (BCA) Protein Assay Kit (Thermo Fisher Scientific, Cat# 23227). Protein samples were then supplemented with 5% 2-mercaptoethanol followed by protein denaturation at 95 °C for 10 min before loaded on the gel. For testing mitochondrial oxidative phosphorylation (OXPHOS) complex subunits, protein samples were incubated at 37 °C for 30 min instead. Equal amounts of proteins were subject to 4-15% Mini-PROTEAN precast gels (Bio-Rad, Cat# 4568084) in 1x running buffer (Bio-Rad, Cat# 1610732) and then transferred onto polyvinylidene difluoride (PVDF) membranes (Bio-Rad, Cat# 1620177) in transfer buffer (3.5 g Tris-base, 15 g glycine, and 200 mL methanol in 1 L) using a transblot apparatus (Bio-Rad, Hercules, CA) at 100 V for 2 hours. The PVDF membranes were then stained with Ponceau S solution (Thermo Fisher Scientific, Cat# A40000279) to assess protein transfer efficiency and total protein loading. The membranes were blocked with 5% milk in Tris-buffered saline with 0.2% Tween 20 (TBST) at room temperature for 1 hour to prevent antibodies from non-specific binding and then incubated with primary antibodies (CD36, 1:500, Thermo Fisher Scientific, Cat# PA1-16813; CHD4, 1:10000, EpiCypher, Cat# 13-2016 (for Figure 1B); CHD4, 1:1000, Abcam, Cat# ab72418 (for Figure 5B); NFIA, 1:1000, Sigma-Aldrich, Cat# HPA006111; OXPHOS antibody cocktail, 1:250, Thermo Fisher Scientific, Cat# 45-8099) overnight at 4 °C. After washing off unbound primary antibodies using TBST three times, the membranes were incubated with the corresponding horseradish peroxidase (HRP) conjugated goat anti-mouse IgG (Jackson ImmunoResearch, Cat# 115-035-174) or goat anti-rabbit IgG (Jackson ImmunoResearch, Cat# 211-032-171) secondary antibodies (1:1000) at room temperature for 1 hour. After washing off unbound secondary antibodies using TBST three times, the membranes were incubated with Pierce^TM^ECL Plus Western Blotting Substrate (Thermo Fisher Scientific, Cat# 32132) and then ready for imaging using OPTIMAX X-Ray Film Processor.

**Figure 1.**
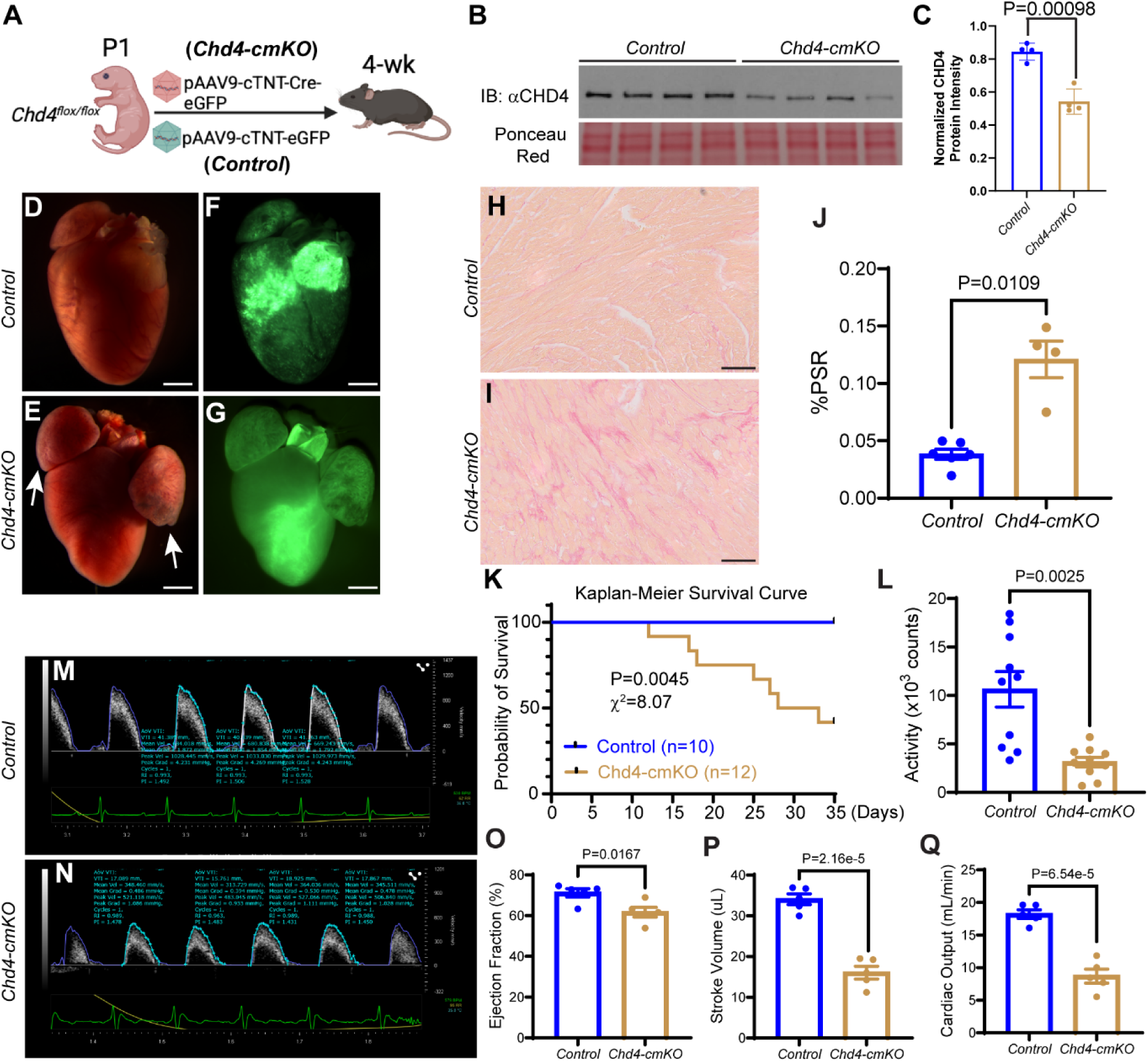
*Chd4* deficiency in postnatal cardiomyocytes results in cardiac dysfunction. (A) Workflow of Adeno-associated viral (AAV) particles-mediated cardiac *Chd4* knockdown in neonates. *Chd4^flox/flox^* neonates at postnatal day 1 (P1) was subcutaneously injected with pAAV9-cTNT2-Cre-eGFP (*Chd4-cmKO*) or pAAV9-cTNT2-eGFP (littermate *Control*) viral particles. Mouse cardiac function, histology, and transcriptome were evaluated at four weeks post viral treatment. (B) Western blot of the whole heart lysate followed by probing anti-CHD4 antibody. Before antibody probing, the blot was stained with Ponceau S Red solution to present the protein loading amount in each lane. (C) Quantification of the CHD4 protein band intensity in (B), normalized to Ponceau S Red loading control. P value: unpaired Student’s t test. (D-G) Representative images of the *Control* (D and F) and *Chd4-cmKO* (E and G) mouse hearts at four weeks post viral treatment. D and E: bright field. F and G: GFP view. White arrows: enlarged atria. Scale bars: 1 mm. (H, I) Representative images of the Picrosirius red (PSR)-stained heart sections. Scale bars: 50 μm. (J) Quantifications of the percent of fibrotic tissue, calculated by dividing the fibrotic tissue pixels by the total tissue pixels. P value: unpaired Student’s t test. (K) Kaplan-Meier curve of the *Chd4-cmKO* (n=12) and *Control* (n=10) mice over five weeks post viral treatment (four litters in total). Mice harvested at four weeks for biochemical analyses were from separate litters/treatment and were not included in this survival quantification. P value and χ^2^: Log-rank test. (L) Scurry Activity Wheel Chamber System was used to record voluntary mouse running for 24 hours for the *Chd4-cmKO* and *Control* mice at four weeks. N=10 (five male and five female) per group. P value: unpaired Student’s t test. (M, N) Representative pulsed-wave (PW) Doppler echocardiographic images of the aortic flow (AF) profiles comparing the peak velocity and velocity-time integral (VTI) of a *Control* mouse (M) and a *Chd4-cmKO* (N) mouse at four weeks post viral treatment. (O-Q) Quantifications of ejection fraction (EF, %), stroke volume (mL), and cardiac output (CO, mL/min) from the echocardiograms. N=5 mice per group. P value: unpaired Student’s t test.

### *In-situ* proximity ligation assay (PLA)

PLA was performed as previously described^20^. Briefly, isolated cardiomyocytes were seeded onto circular coverslips and cultured for 48 hours. The cells were then fixed and permeabilized using 4% paraformaldehyde and 1% Tween-20 in PBS for 10 min and subsequently blocked with a blocking buffer consisting of 10% normal goat serum and 1% Triton X-100 in PBS for 1 hour at room temperature. The cells were then incubated with two specific primary antibodies in a humid chamber at 4 °C overnight: mouse anti-CHD4 (1:250; Sigma-Aldrich, Cat# MABE455) and rabbit anti-NFIA (1:250; Sigma-Aldrich, Cat# HPA006111). Anti-CHD4 and anti-IgG antibodies were co-incubated as the control group. After washing the cells twice with Wash Buffer A for 5 min each time, the cells were then sequentially incubated with PLUS and MINUS PLA probes at 37 °C for 1 hour, with ligase at 37 °C for 30 min, and with polymerase for amplification at 37 °C for 100 min, by using the Duolink® In Situ Red Starter Kit Mouse/Rabbit (Sigma-Aldrich, Cat# DUO92101-1KT). After the final wash with 1% Wash Buffer B for 1 min, the cells were then incubated with an anti-tropomyosin antibody (1:50; Developmental Studies Hybridoma Bank, Cat# CH1) for 1 hour at room temperature, followed by incubation with goat anti-mouse Alexa Fluor 488-conjugated secondary antibody (1:1000; Thermo Fisher Scientific, Cat# A-11001) for 1 h at room temperature. The slides were then mounted using the PLA mounting medium containing DAPI. Imaging was conducted using a Zeiss LSM 880 laser-scanning confocal microscope with a 63× oil objective.

### Statistical methods

The GraphPad Prism (Version 11.0.2; GraphPad Software, Inc., La Jolla, CA) was primarily used for statistical analyses. All data was examined for normality using the Shapiro–Wilk test and for homogeneity of variance using an F-test before parametric statistical analyses were applied. All continuous variables were presented as scatter dot plots with Mean±SEM superimposed unless indicated otherwise. Differences between two groups were evaluated using a two-tailed, unpaired Student’s *t*-test for parametric data, with Welch’s correction applied when the assumption of equal variances was not met. When there were two different grouping factors, two-way ANOVA followed by Tukey’s multiple comparisons test were applied. Log-rank test was used for Kaplan-Meier survival analysis. Benjamini-Hochberg procedure (FDR) followed by multiple comparisons adjustment was used for analyzing RNA-seq data. A *p*-value or adjusted *p*-value <0.05 was considered statistically significant.

## RESULTS

### *Chd4* deficiency in postnatal cardiomyocytes results in cardiac dysfunction and mortality

Previous studies using *Myh6-Cre* and *Corin-Cre* drivers to delete *Chd4* specifically in cardiomyocytes perinatally showed that mutant mice developed normally but died suddenly at weaning age or by 3 months, respectively^17^. To more specifically focus on the postnatal stage, we used an adeno-associated virus (AAV serotype 9)-mediated cardiac Troponin T (cTNT) *Cre* recombinase^37,43,44^ (co-expressed with an eGFP reporter, pAAV9-cTNT-Cre-eGFP) to delete *Chd4* in cardiomyocytes of *Chd4^flox/flox^*mice (designated as *Chd4-cmKO* hereafter, with pups treated with pAAV-cTNT-eGFP serving as the *Control* group) at postnatal day 1 (P1) (2×10^11^ viral genome/pup). Four weeks post-AAV injection, cardiac function, morphology, histology, and transcriptome profiles were evaluated (Figure 1A). The specificity and efficiency of CHD4 protein depletion in cardiomyocytes were confirmed by immunohistochemistry staining and western blotting (Figures 1B, C and S1A). At four weeks, *Chd4-cmKO* mouse body weight was slightly but significantly lower than that of *Control* mice (Figure S1B), and approximately 30% of *Chd4-cmKO* hearts displayed enlarged atria (Figure 1D-G) and exhibited increased interstitial fibrosis in the ventricular myocardium (Figure 1H-J), consistent with previous findings by Gomez-del Arco et al^17^. In a separate cohort monitored over a five-week period, *Chd4-cmKO* mice began succumbing as early as 14 days post-viral treatment. By four weeks, ∼50% of *Chd4-cmKO* mice had died, and only ∼40% survived to five weeks post-viral treatment (P = 0.0045, Log-rank test) (Figure 1K). Together, these findings demonstrate that postnatal cardiomyocyte-specific *Chd4* deletion causes severe cardiac dysfunction, pathological remodeling, and mortality.

To determine whether physical activity was impacted by *Chd4* inactivation in cardiomyocytes, voluntary running was recorded for a continuous 24-hour session using the Scurry Activity Wheel Chamber System with free access to water and diet^45^. *Chd4-cmKO* mice exhibited ∼60% lower activity (Figure 1L) with no significant difference in diet uptake (Figure S1C). To test if reduced exercise tolerance correlated with compromised cardiac systolic function, we performed echocardiography. M-mode and Doppler analyses (Figures 1M, N and S1D-G) revealed that *Chd4-cmKO* mice had reduced systolic function, including reduced ejection fraction (61.70 ± 2.43 vs. 71.23 ± 2.03) (Figure 1O) and fractional shortening (31.88 ± 1.69 vs. 39.68 ± 1.58) (Figure S1H) compared with *Control* mice. Consistent with impaired cardiac function, *Chd4-cmKO* mice displayed profound decreases in stroke volume (16.05 ± 1.56 vs. 34.06 ± 1.32) (Figure 1P), cardiac output (8.71 ± 1.07 vs. 18.22 ± 0.66) (Figure 1Q), and left ventricular end-diastolic diameter (2.65 ± 0.12 vs. 3.41 ± 0.07) (Figure S1I). However, heart rate (Figure S1J) and normalized left ventricular mass (Figure S1K) remained comparable between groups (P > 0.05). These echocardiographic analyses reveal CHD4 supports normal postnatal cardiac function.

### *Chd4* deficiency in postnatal cardiomyocytes dysregulates fatty acid metabolic pathways

To investigate the transcriptomic consequences of cardiomyocyte-specific *Chd4* deletion, bulk RNA-seq analysis was performed with ventricular tissues from *Chd4-cmKO* and *Control* mice. Principal component analysis (PCA) revealed pronounced segregation of the two groups along the primary axis (PC1), which accounted for 88% of the total variance and reflected a highly consistent transcriptomic distinction upon loss of CHD4 (Figure S2A). Differentially expressed gene (DEG) analysis identified 2,662 upregulated genes and 1,702 downregulated genes in *Chd4-cmKO* hearts (Cutoffs for these DEGs: |Log_2_(Fold Change) > 0.5| and adjusted P value < 0.05). Hierarchical clustering confirmed the robust separation (Figure 2A). Notably, downregulated DEGs in *Chd4-cmKO* hearts included key transcriptional regulators and metabolic genes associated with postnatal cardiac maturation, including *Ppara* and *Ppargc1a*, which regulate fatty acid oxidation and mitochondrial biogenesis and function, respectively^46,47^, and *Cpt2*, *Cd36*, *Fabp3*, *Lpl*, and *Slc27a1*, which are involved in fatty acid utilization and transport^48–56^. *Plin5*, a lipid droplet-coating protein involved in cardiac lipid storage and utilization^57,58^, was also downregulated (Figure 2A, blue). Conversely, upregulated DEGs included stress-responsive and fetal cardiac markers such as *Nppa* and *Nppb*, glycolytic enzymes *Eno1* and *Eno2*, and cell cycle regulators, including *Ccne1*, *Ccnb1*, and *Aurkb*^1^ (Figure 2A, red). Analysis of sarcomeric genes further showed a pathological reversion toward immature contractile isoforms, characterized by the downregulation of adult-enriched isoforms (*Myh6, Tnni3, Myl2*) and concurrent retention or reactivation of their corresponding embryonic isoforms (*Myh7, Tnni1, Myl7*)^1,44^ (Figure S2B). Consistently, Gene Set Enrichment Analysis (GSEA) revealed that downregulated genes were significantly enriched in mature metabolic pathways, including oxidative phosphorylation, fatty acid metabolism, and lipid handling (adipogenesis hallmark), whereas upregulated genes were enriched in developmental structural programs (myogenesis), epithelial-mesenchymal transition (EMT), and apical junction pathways (Figures 2B and S2C). Together, these transcriptomic signatures indicate that CHD4 deficiency disrupts the expression of gene programs associated with postnatal metabolic and structural maturation while promoting the reactivation of selected fetal gene programs.

**Figure 2.**
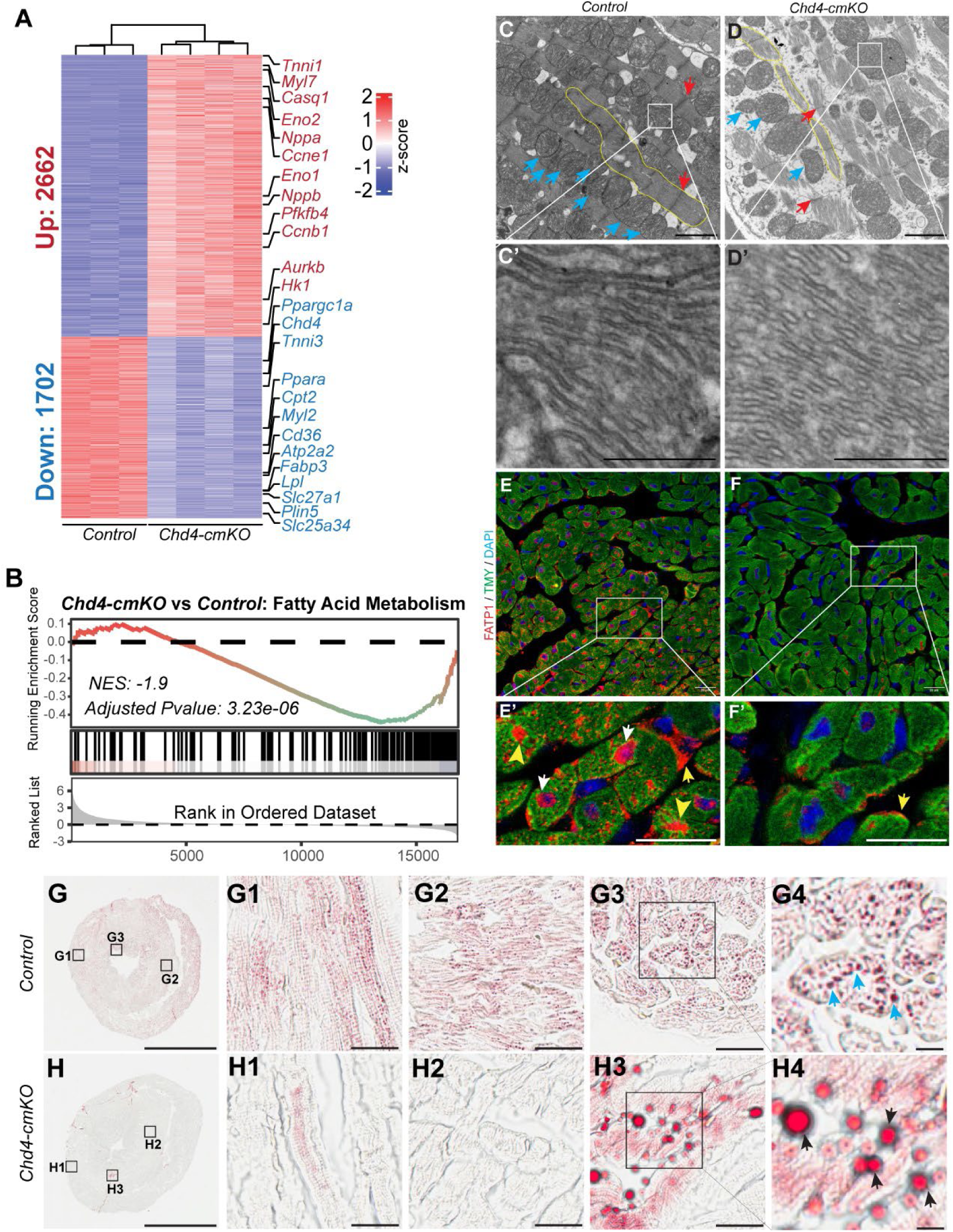
*Chd4* deficiency in postnatal cardiomyocytes dysregulates fatty acid metabolic pathways. (A) Heatmap of differentially expressed genes (DEGs). Whole hearts of the *Chd4-cmKO* and *Control* mice at four weeks post viral treatment were harvested and subjected for RNA isolation and bulk RNA-sequencing. Cutoffs for DEGs: | Log_2_(Fold Change) | > 0.5 and adjusted P value < 0.05. Color bar: z-score. (B) Gene Set Enrichment Analysis (GSEA) plot of the genes involved in the fatty acid metabolic pathway. NES: normalized enrichment score. (C, D) Representative transmission electron microscopy images of the *Control* and *Chd4-cmKO* mouse hearts at four weeks post viral treatment. Yellow area: Myofibrils. Blue arrows: Mitochondria. Red arrows: Z-disc. Scale bars: 2 μm (C and D), 500 nm (C’ and D’). (E, F) Representative immunohistochemistry-stained images of the *Control* and *Chd4-cmKO* heart sections. Red: FATP1 (arrows); green: Tropomyosin (TMY), marker of cardiomyocytes; blue: DAPI. Scale bars: 20 μm (E and F), 60 μm (E’ and F’). (G, H) Representative Oil Red O-stained images of the *Control* and *Chd4-cmKO* heart sections. G1-G4 and H1-H4: higher magnification of outer layer of left ventricle (G1 and H1), ventricular septum (G2 and H2), and inner layer of left ventricle (G3 and H3). Scale bars: 2 mm (G and H), 30 μm (G1-G3 and H1-H3), 6 μm (G4 and H4).

We next employed transmission electron microscopy (TEM) to determine whether ultrastructural morphology aligned with these transcriptomic alterations. *Control* hearts exhibited well-organized myofibrils assembled into individual contractile units anchored by clear Z-discs (Figure 2C, red arrows). In contrast, *Chd4-cmKO* hearts displayed a severe reduction in myofilament density accompanied by the loss or deformation of Z-discs (Figure 2D, red arrows). We also observed clear mitochondrial abnormalities in *Chd4-cmKO* hearts, including decreased mitochondrial density and fragmented mitochondrial cristae (Figure 2D, D’) compared with the continuous, linear cristae observed in *Controls* (Figure 2C, C’). These defective myofibrillar and mitochondrial structures, combined with the transcriptomic downregulation of oxidative phosphorylation and fatty acid metabolic genes, indicate that CHD4 is required for postnatal metabolic maturation. We next examined the protein expression of two key fatty acid transporters, CD36 (encoded by *Cd36*) and fatty acid transport protein 1 (FATP1, encoded by *Slc27a1*), both of which contribute to myocardial fatty acid uptake^50,51,54,55,59^. Consistent with the downregulation of *Cd36* and *Slc27a1* transcripts, immunostaining revealed reduced expression of both transporters in *Chd4-cmKO* hearts. Consistent with previously reported FATP1 cellular localization^59^, FATP1 was detected predominantly at the cardiomyocyte surface, in the cytoplasm, and in perinuclear regions in *Control* hearts (Figure 2E, E′). In contrast, FATP1 staining was markedly diminished in *Chd4-cmKO* hearts, with minimal signal detected at the cell surface or in the cytoplasm (Figure 2F, F′). Similarly, CD36 was robustly expressed at the cardiomyocyte surface in *Control* hearts, whereas its expression was reduced in *Chd4-cmKO* hearts (Figure S2D-G).

Because transcriptomic programs for fatty acid handling and oxidative pathways were suppressed, we next evaluated myocardial lipid accumulation. Oil Red O staining revealed neutral lipid accumulation within cardiomyocytes in *Control* hearts (Figure 2G–G4), whereas lipid-positive droplets in *Chd4-cmKO* hearts were predominantly detected in the myocardial interstitial spaces (Figure 2H-H4). Oroboros high-resolution respirometry further demonstrated impaired mitochondrial respiratory capacity in *Chd4-cmKO* hearts, characterized by reduced maximal electron transfer system (ETS) capacity and Complex IV (CIV)-supported respiration (Figure S2H). In contrast, absolute FAO-supported oxygen consumption was not significantly different between genotypes, despite the suppression of FAO-related gene expression. Notably, flux control ratios (FCRs) for FAO and several substrate-supported respiratory states were increased in mutant hearts (Figure S2I), likely reflecting the reduced maximal respiratory capacity rather than enhanced absolute FAO capacity. Consistent with these functional defects, immunoblot analysis revealed altered abundance of representative subunits of mitochondrial oxidative phosphorylation complexes in *Chd4-cmKO* hearts (Figure S2J). Collectively, these findings indicate that CHD4 is required to maintain the postnatal fatty acid metabolic gene program, normal myocardial lipid distribution, and mitochondrial respiratory capacity.

### CHD4 regulates fatty acid metabolic genes specifically in the postnatal heart

Because metabolic maturation occurs primarily after birth and CHD4 functions in both embryonic and postnatal hearts, we next asked whether CHD4-dependent regulation of fatty acid metabolic genes is specific to the postnatal stage. To address this question, we integrated six independent transcriptomic datasets of cardiac *Chd4* mutants, including five previously published datasets^17,22^ and the new postnatal dataset generated in this study (Figure 2A). These datasets comprised three embryonic and three postnatal transcriptomic studies. By comparing differentially expressed genes (DEGs) across datasets, we sought to identify stage-specific CHD4-regulated gene programs, particularly those involved in fatty acid metabolism. We defined CHD4-mediated dysregulated genes (DysGenes) as DEGs in at least two independent datasets within each developmental stage to minimize potential biases arising from differences in *Cre* drivers, sequencing platforms, and bioinformatic pipelines (Figure 3A–D). This cross-dataset analysis identified 71 DysGenes downregulated in embryonic cardiac *Chd4* mutants (Figure 3A), 232 downregulated in postnatal cardiac *Chd4* mutants (Figure 3B), 396 upregulated in embryonic cardiac *Chd4* mutants (Figure 3C), and 586 upregulated in postnatal cardiac *Chd4* mutants (Figure 3D). The statistical significance of overlaps between gene lists was assessed using the hypergeometric test, with a mouse background genome size of 20,000 genes. We next intersected these four DysGene lists and identified 472 and 214 DysGenes uniquely up-and down-regulated, respectively, in postnatal cardiac *Chd4* mutant hearts (Figure 3E). Importantly, these 214 genes, including *Slc27a1*, *Lpl*, *Fabp3*, *Nudt8*, *Ndufa8*, and *Cox7a1*, were significantly enriched in biological processes related to fatty acid metabolism, lipid catabolism, oxidative phosphorylation, and acyl-CoA metabolism. Notably, many of these genes encode mitochondrial proteins involved in mitochondrial biogenesis and function, including *Mtfp1* and *Mtg1* (Figure 3F). We also identified 112 DysGenes upregulated in both embryonic and postnatal cardiac *Chd4* mutant hearts, which were significantly enriched in skeletal muscle and neuromuscular gene programs (including *Tnni2, Tnnc2, Tnnt3, Camk2b*, and *Ache*), aligning with CHD4’s typical transcriptional repressor function^17,22^. By contrast, only 14 downregulated DysGenes were shared between embryonic and postnatal cardiac *Chd4* mutant hearts and were not enriched in any specific gene programs (Figure 3E). These results indicate that CHD4 coordinates distinct transcriptional programs in embryonic versus postnatal hearts.

**Figure 3.**
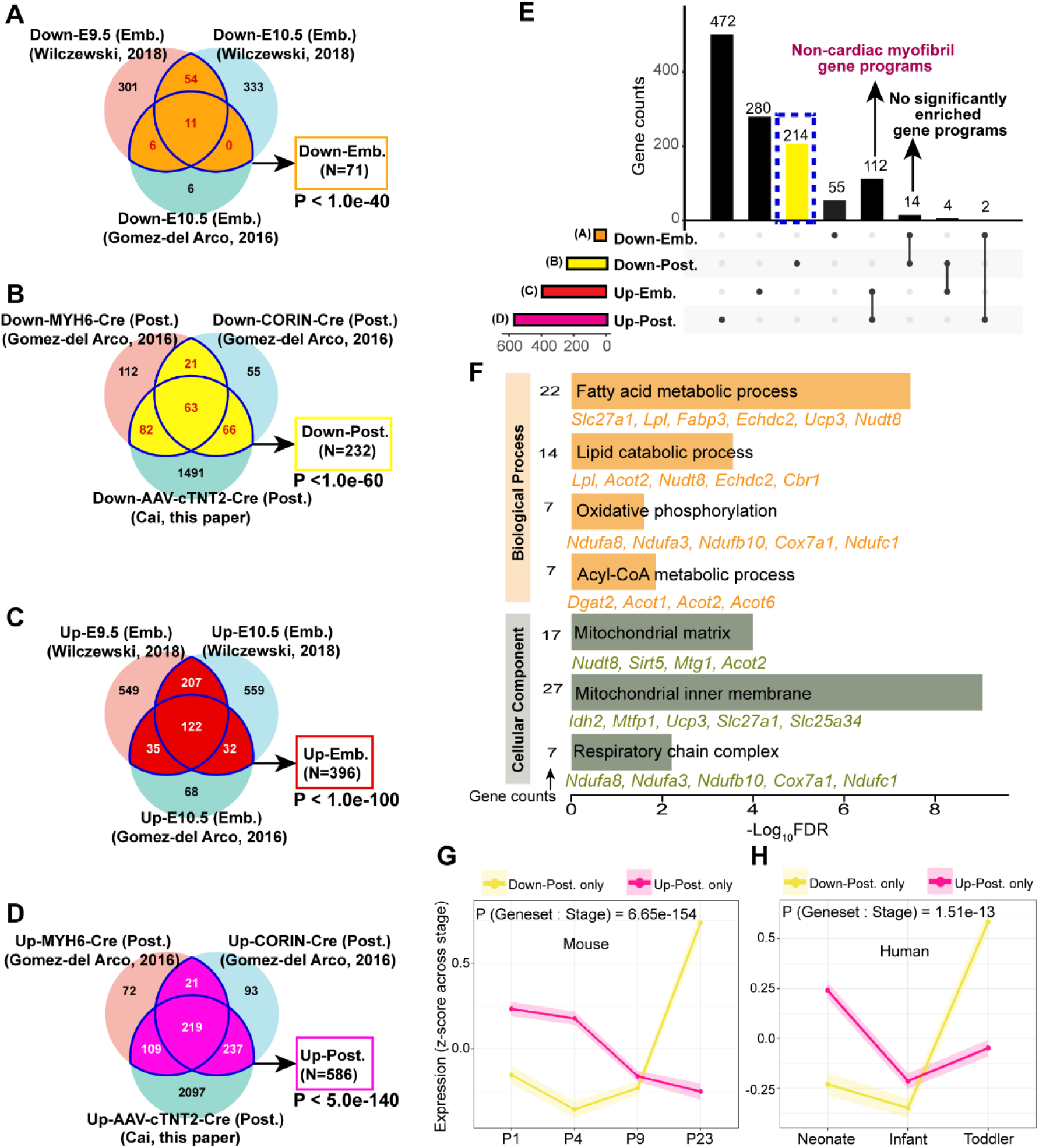
CHD4 regulates fatty acid metabolic genes specifically in the postnatal heart. (A) Venn diagrams of downregulated DEGs in embryonic *Chd4-cmKO* mouse hearts. (B) Venn diagrams of downregulated DEGs in postnatal *Chd4-cmKO* mouse hearts. (C) Venn diagrams of upregulated DEGs in embryonic *Chd4-cmKO* mouse hearts. (D) Venn diagrams of upregulated DEGs in postnatal *Chd4-cmKO* mouse hearts. In A-D, genes existing in at least two datasets were defined as dysregulated genes (DysGenes) in each overlapping. Statistical significance of the DysGenes across datasets was calculated using a one-tailed Hypergeometric test against a standard background mouse genome size. Embryonic *Chd4-cmKO* mouse heart DEGs and *Myh6Cre*-or *CorinCre*-mediated postnatal *Chd4-cmKO* mouse heart DEGs were retrieved from Wilczewski et al, 2018 and Gomez-del Arco et al, 2016. AAV9-cTNT2-Cre-eGFP-mediated postnatal *Chd4-cmKO* DEGs were generated in this paper (heatmap of which is shown in Fig. 2A). (E) Upset plot showing overlaps of DysGenes in A-D. (F) Overrepresented Biological Process and Cellular Component GO (gene ontology) terms of the 214 “Down-Post. only” genes, with gene counts and representative genes in each term labeled. FDR: false discovery rate. (G, H) Genes only upregulated in postnatal *Chd4-cmKO* mouse hearts (“Up-Post. only”, N=472 genes) and genes only downregulated in postnatal *Chd4-cmKO* mouse hearts (“Down-Post. only”, N=214 genes) (from Fig. 3E) were overlapped to a wildtype postnatal mouse heart transcriptomic dataset across four stages: postnatal day 1 (P1), P4, P9, and P23 (Talman et al., 2018) and to a human heart transcriptomic dataset of three postnatal stages: neonate, infant, and toddler (BioStudies database (http://www.ebi.ac.uk/biostudies) under accession number E-MTAB-6814). 445 of the 472 “Up-Post. only” genes and 196 of the 214 “Down-Post. only” genes were found in the Talman mouse dataset (G) and 401 of the 472 “Up-Post. only” genes and 171 of the 214 “Down-Post. only” genes were found in the BioStudies human dataset (H). For each overlapped gene, TPM (Transcripts Per Million) values at each stage were averaged and z-scored across that gene’s own stages. Lines show mean z-scored expression per gene set (“214 Down-Post. only”-gene, yellow; “472 Up-Post. only”-gene, pink) at each stage, with shaded ribbons indicating ± SEM across genes. Whether trajectory shape differed between gene sets was tested with a linear mixed model (log2(TPM+1) ∼ Geneset × Stage + (1 | Gene)); the reported P value is for the Geneset:Stage interaction term.

Given the enrichment of the 214 postnatally downregulated DysGenes in metabolic pathways, we next examined their normal expression trajectories during early postnatal heart development in mice and humans. We mapped these 214 DysGenes to a wild-type postnatal mouse heart transcriptomic dataset across four stages (postnatal days P1, P4, P9, and P23)^60^ and to a human heart transcriptomic dataset spanning three postnatal stages (neonate, infant, and toddler; BioStudies accession E-MTAB-6814) and identified 196 and 171 matching genes in the mouse and human datasets, respectively (Figure S3A, B). For comparison, we similarly mapped the 472 upregulated DysGenes (Figure 3E) and identified 445 and 401 matching genes in mice and humans, respectively (Figure S3A, B). For each mapped gene, Transcripts Per Million (TPM) values from corresponding studies were averaged and z-scored across stages. Strikingly, the DysGenes downregulated in postnatal cardiac *Chd4* mutant hearts showed a significant increase from early neonatal stages (P1/P4 in mice; neonates in humans) to late stages (P23 in mice; toddlers in humans). Conversely, DysGenes upregulated in postnatal cardiac *Chd4* mutant hearts showed a significant decrease from early to late stages (Figure 3G, H). Examples of genes exhibiting the developmental upregulation pattern included *Mb*, encoding myoglobin, which facilitates oxygen storage and delivery in muscle^61^; *Rpl3l*, encoding a ribosomal protein enriched in striated muscle^62,63^; and *Ndufa8*, encoding a subunit of mitochondrial respiratory complex I (Figure S3C–H)^64^. Together, these findings reveal that genes downregulated in postnatal cardiac *Chd4* mutant hearts normally increase in expression during postnatal development in both mice and humans, coinciding with the establishment of mature cardiac metabolic programs.

### Developmental changes in CHD4 chromatin occupancy in the postnatal heart

Given that these 214 DysGenes are selectively downregulated in postnatal *Chd4* mutant hearts, we next investigated whether CHD4 directly binds these genes and whether its binding patterns differ between embryonic and postnatal hearts^21,22^. Integrating CHD4 ChIP-seq datasets from embryonic and postnatal wild-type mouse hearts revealed that 36 of these genes lacked CHD4 binding at any stage. In contrast, 102 genes were bound by CHD4 in both stages, and 67 genes were bound exclusively in the postnatal heart, and only 9 were bound exclusively in the embryonic heart (Figure 4A). Overall, 111 genes were bound by CHD4 in embryonic hearts and were associated with 200 peaks (∼1.8 peaks/gene), whereas 169 genes were bound in postnatal hearts and were associated with 389 peaks (∼2.3 peaks/gene). These findings indicate more extensive CHD4 occupancy of the analyzed genes in postnatal hearts. To determine whether CHD4 occupied the same genomic regions at both stages among the 102 shared genes, we compared peak distributions and identified more peaks in postnatal than embryonic hearts (249 versus 185). Peak intersection analysis revealed that 104 postnatal peaks overlapped with 134 embryonic peaks, with some broad postnatal peaks spanning multiple narrower embryonic peaks, whereas 145 postnatal peaks had no overlap with embryonic peaks (Figure 4A). These findings reveal extensive developmental differences in CHD4 occupancy, including postnatal-specific binding regions.

**Figure 4.**
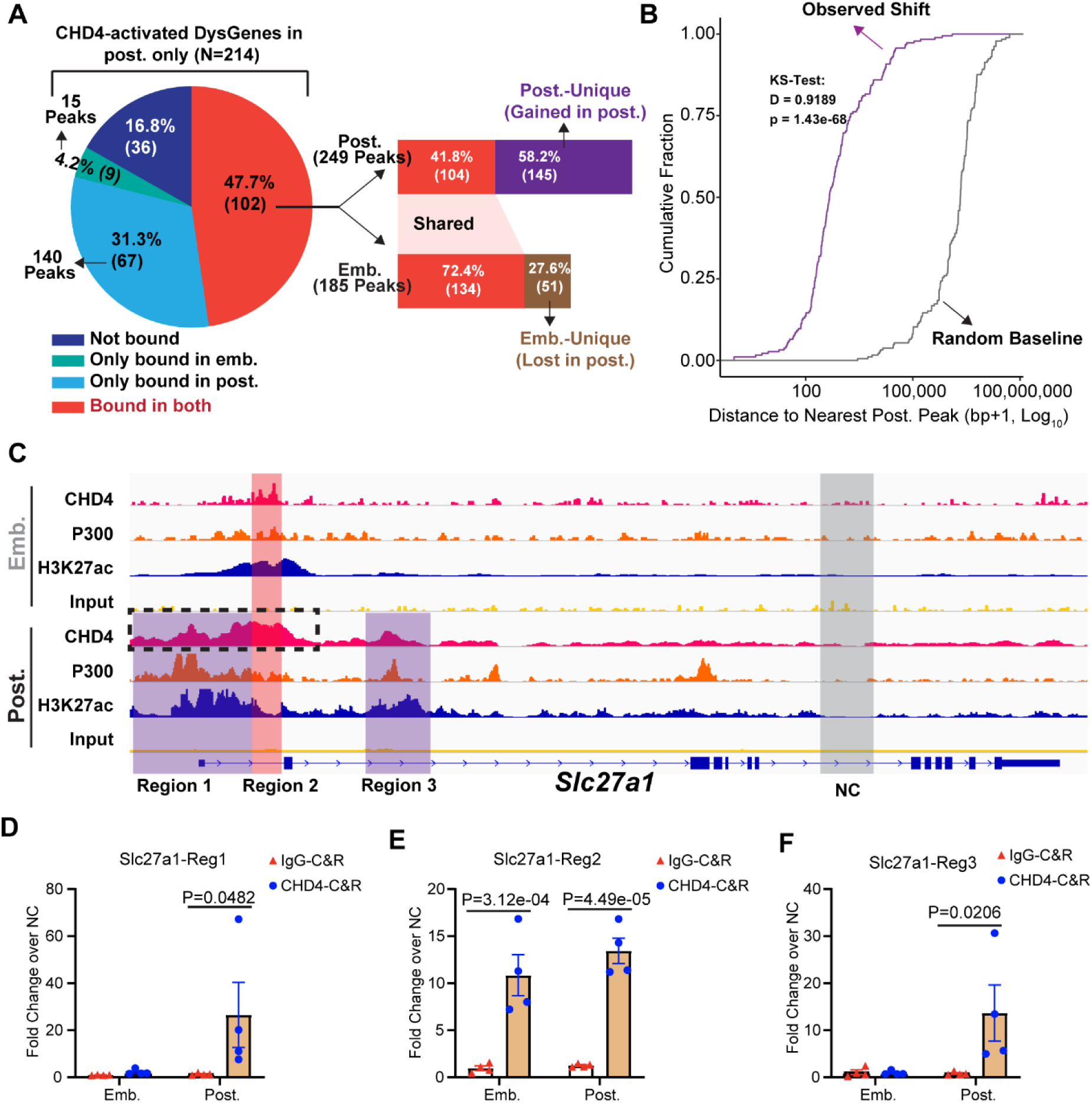
CHD4-bound regions emerging in the postnatal heart. (A) Proportional analysis of CHD4 binding among the 214 “Down-Post. only” genes. Left: Number of genes bound by CHD4 in the embryonic and postnatal hearts. Right: Genomic binding analysis for the genes (N=102) bound by CHD4 in both embryonic and postnatal hearts, highlighting 145 regions uniquely detected in postnatal hearts. (B) Empirical Cumulative Distribution Function (ECDF) showing the shortest distances from embryonic CHD4 peak centers to the nearest postnatal binding site (Observed, purple curve) versus a permuted random genomic baseline (Random Baseline, grey curve). The pronounced leftward shift of the observed curve indicates that CHD4 binding turnover occurs via local neighborhood reallocation rather than random genomic redistribution (D = 0.9189, P = 1.43e-68, one-sided Kolmogorov-Smirnov test). (C) Representative Integrative Genomic Viewer (IGV) browser tracks of the CHD4-bound regions on the *Slc27a1* gene. Region 2 (highlighted in red) is bound by CHD4 in both embryonic and postnatal hearts, whereas Regions 1 and 3 represent postnatal binding sites (highlighted in purple). A Region with few reads in both embryonic and postnatal CHD4-ChIP-seq tracks were selected as the negative control (NC) region (highlighted in grey) used for CHD4-CUT&RUN (Cleavage Under Targets & Release Using Nuclease) enrichment calculation. ChIP-seq tracks of the active histone mark H3K27ac and the histone acetyltransferase P300 in embryonic and postnatal mouse hearts were used to indicate the active transcriptional feature of highlighted regions. ChIP-seq data were retrieved from literature. CHD4-ChIP-seq in the embryonic mouse heart (E10.5): Wilczewski et al, 2018 (GSE109012). CHD4-ChIP-seq in the postnatal mouse heart (atria) (P24): Sweat et al., 2025 (GSE284095). P300-ChIP-seq in the embryonic (E12.5) and postnatal (P28) mouse heart: Zhou et al., 2023 (GSE195901). H3K27ac-ChIP-seq in the embryonic mouse heart (E10.5): Gu et al., 2022 (GSE213572). H3K27ac-ChIP-seq in the postnatal mouse heart (atria) (P28): Sweat et al., 2023 (GSE222970). (D-F) Cardiomyocytes were isolated from wildtype C57Bl/6 mouse hearts at E10.5 or P1-P2 and cultured for 72 hours. CHD4-and IgG-CUT&RUN were performed using the isolated cardiomyocytes followed by library preparation and region-targeted quantitative PCR (qPCR). Genomic DNA purified from isolated embryonic or neonatal cardiomyocytes, without antibody targeting, was used as an input DNA to normalize the enrichment signal and the selected NC Region (indicated in Fig. 4C) was used to calculate the enrichment fold of Regions 1-3. P values: two-way ANOVA.

**Figure 5.**
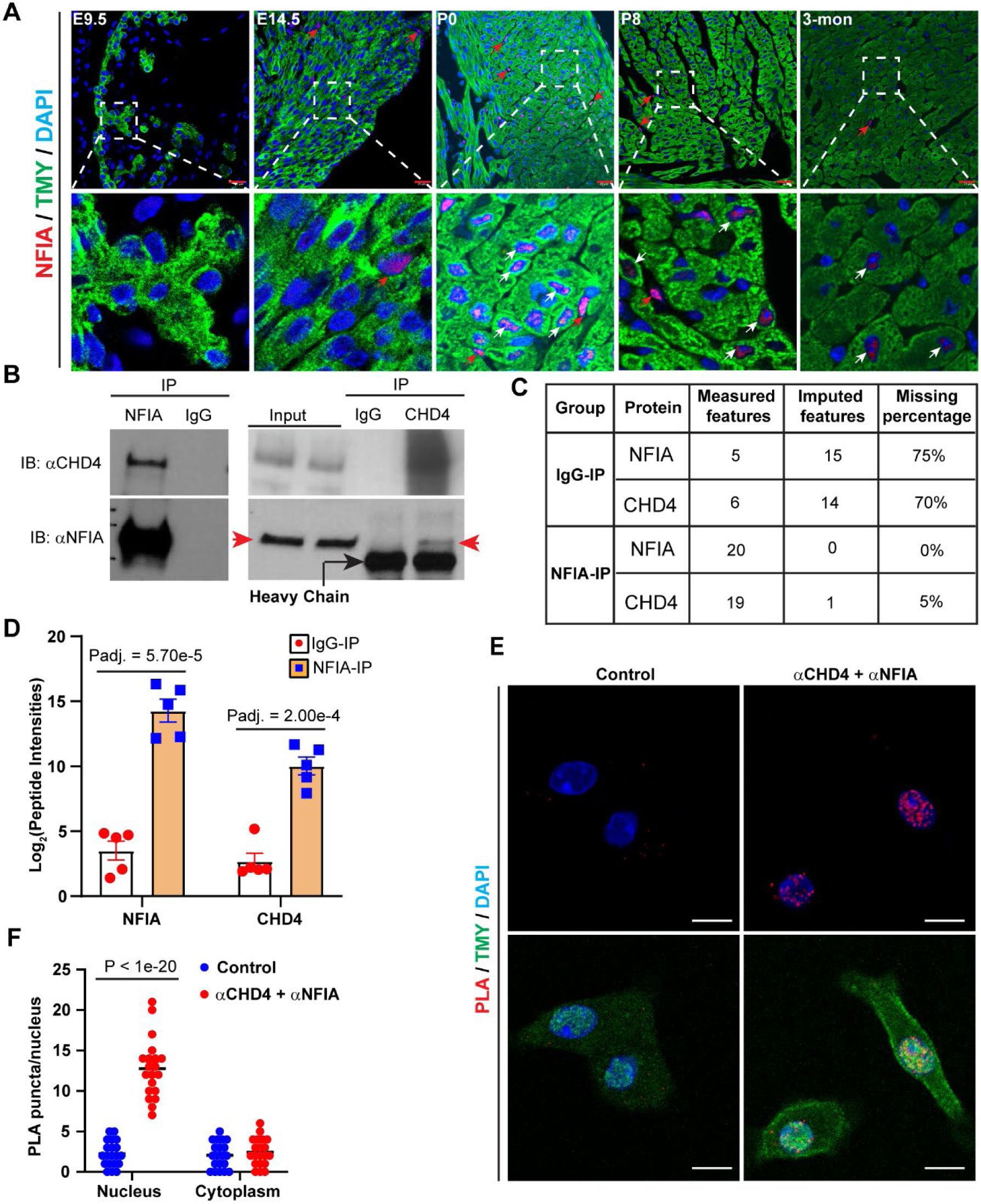
CHD4 interacts with NFIA in postnatal cardiomyocytes. (A) Representative immunohistochemistry-stained images of the wildtype C57Bl/6 mouse heart sections at indicated stages. Red: NFIA (arrows); green: Tropomyosin (TMY), marker of cardiomyocytes; blue: DAPI. Red arrows: NFIA^+^ noncardiomyocytes. White arrows: NFIA^+^ cardiomyocytes. N=3 individual hearts per stage. Scale bars: 20 μm. (B) Immunoprecipitation (IP) with anti-IgG (control), anti-NFIA, or anti-CHD4 antibodies on isolated cardiac nuclei from P1-P2 wildtype C57Bl/6 mouse hearts followed by western blotting. 5% lysate without adding antibodies was used as input controls. Red arrows: NFIA bands. Black arrow: heavy chain. (C) NFIA-IP was performed on isolated and purified cardiac nuclei from P1-P2 wildtype C57Bl/6 mouse hearts followed by Liquid Chromatography-Tandem Mass Spectrometry (LC-MS/MS) analysis. The numbers of measured and imputed features and the percentage of missing measurements are shown for NFIA and CHD4 in IgG control and NFIA-IP groups. Missing values were imputed during data processing. Values represent the number of measured and imputed features and the percentage of missing measurements for each protein in the indicated group. N=5 biological replicates per IP group. (D) NFIA and CHD4 peptide intensity in the IgG-IP and NFIA-IP samples. (E) Proximity Ligation Assay (PLA) was performed on isolated wildtype P1 cardiomyocytes that were cultured on coverslips. In the PLA assay, anti-CHD4 and anti-NFIA antibodies were co-incubated to identify CHD4-NFIA interaction, while anti-CHD4 and anti-IgG antibodies were co-incubated as the control group. After the PLA assay amplification step, coverslips were co-stained with anti-Tropomyosin (TMY) to label cardiomyocytes. Scale bars: 10 μm. (F) Quantification of PLA puncta in the nucleus and cytoplasm. P value: two-way ANOVA.

To determine whether developmental changes in CHD4 occupancy reflect local redistribution of binding sites rather than random genomic redistribution, we calculated the shortest distance from each embryonic CHD4 peak center to the nearest postnatal consensus binding site (Figure 4B). Compared with a permuted random genomic background, embryonic CHD4 peak centers showed a highly nonrandom spatial distribution, with most located in close proximity to postnatal binding sites (Figure 4B; D = 0.9189, P = 1.43e-68, Kolmogorov-Smirnov test). These findings suggest that developmental changes in CHD4 occupancy occur predominantly within established chromatin neighborhoods (Figure 4B).

Genomic feature analysis further revealed that approximately 72% of shared peaks were located within 1 kb of transcription start sites (TSSs). Stage-specific peaks showed distinct distributions: approximately 10% of embryonic-specific peaks were located within 1 kb of TSSs, compared with approximately 23% of postnatal-specific peaks. In contrast, embryonic-specific peaks were more frequently located in introns (approximately 47% versus 36% of postnatal-specific peaks; Figure S4A).

A representative example of these binding dynamics is *Slc27a1*. In embryonic hearts, CHD4 occupied the *Slc27a1* promoter (within 1 kb of the transcription start site) at a narrow peak of approximately 400 bp (Region 2, red). In postnatal hearts, this region overlapped with a broader CHD4 peak of approximately 3,000 bp, most of which represented postnatal-specific occupancy (dashed square; e.g., Region 1, purple). An additional CHD4 peak was detected exclusively in the postnatal heart at a nearby promoter-proximal region (Region 3). ChIP-seq tracks for the active histone mark H3K27ac and the histone acetyltransferase p300 showed active regulatory features at these loci in embryonic and postnatal hearts^65–67^ (Figure 4C). Similar developmental changes in CHD4 occupancy were observed at *Lgr6*, which encodes a G protein-coupled receptor that promotes fatty acid metabolism in the hearts of pressure-overloaded mice^68^, and *Dgat2*, which encodes diacylglycerol acyltransferase 2, an enzyme catalyzing the final step of triglyceride synthesis^69^ (Figure S4B, C).

Because the published ChIP-seq datasets utilized whole heart or atrial cardiac nuclei, we validated the developmental patterns of CHD4 occupancy in primary embryonic and postnatal cardiomyocytes using CUT&RUN. Region-specific qPCR was performed using primers that amplified 100–200 bp fragments within the highlighted regions, along with a negative control (NC) region with minimal CHD4 occupancy. Consistent with the ChIP-seq results, CHD4 was enriched at Region 2 of *Slc27a1* in both embryonic and postnatal cardiomyocytes, whereas enrichment at Regions 1 and 3 was detected exclusively in postnatal cardiomyocytes (Figure 4D–F). Similar stage-dependent occupancy patterns were observed at *Lgr6* and *Dgat2* (Figure S4D–H). Together, these findings validate the developmental changes in CHD4 occupancy in cardiomyocytes and demonstrate postnatal-specific CHD4 binding at regulatory regions of genes involved in fatty acid metabolism.

### CHD4 interacts with NFIA in postnatal cardiomyocytes

To identify transcription factors that interact with CHD4 in postnatal cardiomyocytes and potentially facilitate its recruitment to fatty acid metabolic genes, we compared our previously generated CHD4 interactome datasets from embryonic and postnatal ventricles^18,20^. This analysis identified NFIA as a candidate CHD4-interacting partner detected only in the postnatal cardiac CHD4 interactome (Figure S5A). NFIA belongs to the Nuclear Factor I (NFI) family of transcription factors and is broadly expressed across multiple tissues, including the liver, brain, bone, adipose tissue, and heart^70^. Previous studies have established NFIA as an important regulator of brown adipocyte gene programs^28,30^. However, whether NFIA plays a similar regulatory role in cardiomyocytes, particularly in postnatal cardiac metabolic maturation, remains unknown.

To investigate the role of NFIA in regulating CHD4 function, we first examined its spatiotemporal expression during cardiac development. Immunohistochemistry revealed no detectable NFIA expression at E9.5, occasional nuclear expression in non-cardiomyocytes at E14.5, robust nuclear expression in cardiomyocytes at postnatal day 0 (P0), moderate expression at P8, and markedly reduced expression in adult (3-month-old) hearts (Figure 5A). These findings indicate that cardiomyocyte NFIA expression is highest during the early postnatal period.

To validate our previous proteomics findings, we performed immunoprecipitation (IP) using an anti-CHD4 antibody in neonatal cardiac nuclear extracts, followed by western blotting for NFIA, and reciprocal IP using an anti-NFIA antibody similarly recovered CHD4 (Figure 5B). To further validate this interaction in cardiomyocyte nuclei, we isolated PCM1-positive cardiomyocyte nuclei by magnetic-activated cell sorting (MACS), performed NFIA immunoprecipitation, and analyzed the immunoprecipitates by mass spectrometry (Figure S5B–E). Mass spectrometry detected 20 NFIA peptide features in NFIA immunoprecipitates compared with 5 in IgG controls, as well as 19 CHD4 peptide features compared with 6 in IgG controls (Figure 5C). The corresponding peptide intensities were also significantly higher in NFIA immunoprecipitates than in IgG controls (Figure 5D). Together, these findings provide complementary biochemical and proteomic evidence supporting an interaction between NFIA and CHD4 in neonatal cardiomyocyte nuclei.

Finally, to assess the spatial proximity of CHD4 and NFIA in cardiomyocytes, we performed a proximity ligation assay (PLA) in isolated neonatal cardiomyocytes using anti-CHD4 and anti-NFIA antibodies, with anti-CHD4/anti-IgG serving as a negative control. Cells were subsequently co-stained with anti-tropomyosin (TMY) to identify cardiomyocytes and DAPI to visualize nuclei. Robust PLA puncta were detected predominantly within cardiomyocyte nuclei in the anti-CHD4/anti-NFIA group, whereas minimal signal was observed in the cytoplasm or negative control group (Figure 5E, F). Together with the reciprocal immunoprecipitation and mass spectrometry results, these findings provide convergent evidence supporting an interaction between CHD4 and NFIA in postnatal cardiomyocyte nuclei.

### NFIA and CHD4 regulate a same set of metabolic genes

To determine whether NFIA regulates fatty acid metabolic genes similarly to CHD4 in the postnatal heart, we generated cardiomyocyte-specific *Nfia* knockout mice by crossing *Tnnt2-Cre* mice with the *Nfia^flox/flox^* mice (*Tnnt2^Cre/+^*;*Nfia^flox/flox^*, designated *Nfia-cmKO*), which specifically deletes Exon 2 which encodes the NFIA DNA-binding domain^35^ (Figure S6A). Efficient deletion of *Nfia* Exon 2 and loss of NFIA protein specifically in cardiomyocytes were confirmed by region-specific qPCR and immunohistochemistry staining (Figure S6B–D). *Nfia-cmKO* mice were born at expected Mendelian ratios and survived into adulthood. Echocardiography performed at postnatal day 28 (P28)—matching the age evaluated in *Chd4-cmKO* cohorts—revealed that *Nfia-cmKO* mice exhibited a significant reduction in cardiac output (Figure S6E) and a marginal decrease in heart rate (Figure S6F). Although global left ventricular ejection fraction (LV-EF) and fractional shortening (LV-FS) were preserved at this stage (P > 0.05; Figure S6G), mutant hearts displayed a significant increase in the mitral valve (MV) E/A ratio (Figure S6H) and a trend toward a reduced tricuspid valve (TV) E/A ratio (P = 0.123; Figure S6I).

Given the milder cardiac phenotype of *Nfia-cmKO* mice compared with *Chd4-cmKO* mice, we next investigated the extent to which NFIA contributes to the CHD4-dependent transcriptional program in the postnatal heart. We performed bulk RNA-seq on ventricular tissues from *Nfia^flox/flox^* and *Nfia-cmKO* mice at postnatal day 2 (P2) and confirmed efficient ablation of *Nfia* Exon 2 (Figure S6J). Differential gene expression analysis identified 132 downregulated genes and 66 upregulated genes in *Nfia-cmKO* hearts (∣log2 Fold Change∣>0.5, adjusted P<0.05; Figure 6A). Gene Ontology (GO) analysis of the 132 downregulated genes revealed significant enrichment of biological processes related to fatty acid oxidation, lipid metabolism, and regulation of lipid localization (Figure 6B). These findings indicate that NFIA deficiency in cardiomyocytes disrupts the expression of genes involved in postnatal cardiac fatty acid metabolism.

**Figure 6.**
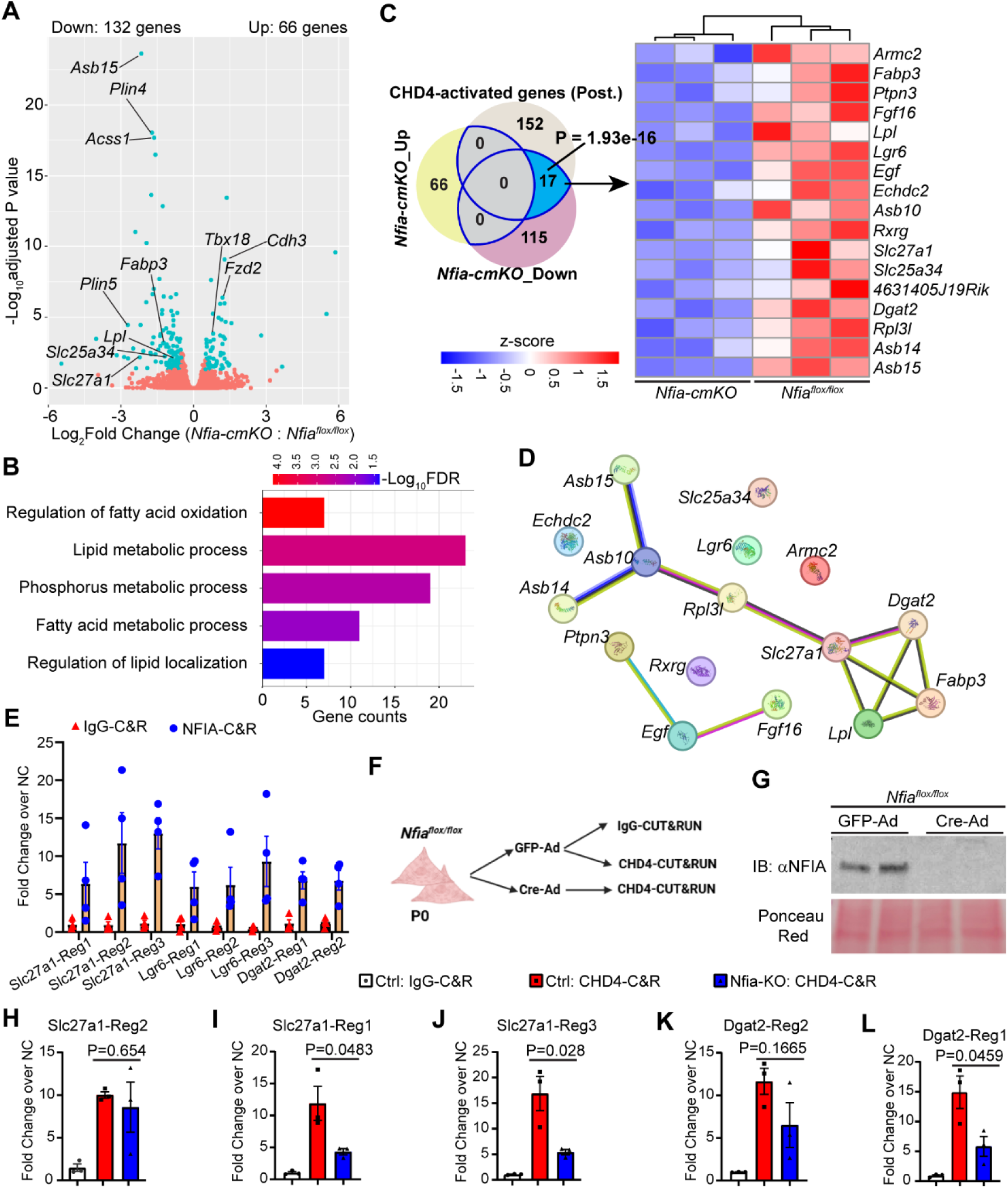
NFIA recruits CHD4 to activate fatty acid metabolic genes in postnatal cardiomyocytes. (A) Volcano plot of the transcriptomes of *Tnnt2^Cre/+^;Nfia^flox/flox^*(*Nfia-cmKO*) and littermate *Nfia^flox/flox^* mouse hearts at P1. Blue dots: differentially expressed genes (DEGs); red dots: non-DEGs. Cutoffs for DEGs: Cutoffs for DEGs: | Log2(Fold Change) | > 0.5 and adjusted P value < 0.05. (B) Overrepresented Biological Process GO terms of the downregulated DEGs in *Nfia-cmKO* mouse hearts. (C) Left: Venn diagram of gene overlapping of the down-and up-regulated DEGs and 169 genes directly bound by and activated by CHD4 (from Fig. 4A). P value: one-tailed Hypergeometric test against a standard background mouse genome size. Right: Heatmap of the 17 overlapped genes in *Nfia-cmKO* and *Nfia^flox/flox^* transcriptomes. Color bar: z-score. (D) Functional protein association networks of the 17 overlapped genes in Fig. 6C, generated with the STRING database. (E) Cardiomyocytes were isolated from wildtype C57Bl/6 mouse hearts at P1-P2 and cultured for 72 hours. NFIA-and IgG-CUT&RUN were performed using the isolated cardiomyocytes followed by library preparation and region-targeted qPCR. Genomic DNA purified from isolated neonatal cardiomyocytes, without antibody targeting, was used as an input DNA to normalize the enrichment signal and selected negative control (NC) Region of each gene was used to calculate the enrichment fold of targeted Regions. (F) Workflow of CHD4-CUT&RUN with *Nfia* knockout neonatal cardiomyocytes. Cardiomyocytes of the *Nfia^flox/flox^*neonates at P1 were isolated and cultured followed by GFP-adenovirus (GFP-Ad; *Control*) or *Cre*-adenovirus (Cre-Ad; *Nfia-KO*) treatment for 72 hours. Then IgG-and CHD4-CUT&RUN were performed. (G) Western blot of the adenoviral particles-treated *Nfia^flox/flox^* cardiomyocyte lysate followed by probing anti-NFIA antibody. Before antibody probing, the blot was stained with Ponceau S Red solution to present the protein loading amount in each lane. (H-L) CUT&RUN region-targeted qPCR of indicated gene regions. P values: unpaired Student’s t test.

We next examined the overlap between the 132 downregulated DEGs identified in *Nfia-cmKO* hearts and the 169 genes bound by CHD4 in the postnatal heart that were previously found to be downregulated following *Chd4* deletion. Intersection analysis identified 17 genes shared between these two gene sets, whereas none of the 66 upregulated DEGs in *Nfia-cmKO* hearts overlapped with the CHD4-bound postnatal gene set. The overlap between the downregulated gene sets was highly significant (P = 1.93 × 10⁻¹⁶, hypergeometric test; Figure 6C). STRING network analysis^71^ further revealed that these 17 shared genes form a functionally interconnected network, featuring a prominent fatty acid metabolism-related cluster, including *Slc27a1*, *Dgat2*, *Fabp3*, and *Lpl*, and an auxiliary subcluster comprising *Asb10*, *Asb14*, and *Asb15*. These three genes encode members of the ankyrin repeat and SOCS box-containing (ASB) protein family, which generally function as substrate-recognition components of E3 ubiquitin ligase complexes involved in protein turnover^72^. This subcluster was connected to the fatty acid metabolism-related cluster through *Rpl3l* (Figure 6D).

### NFIA facilitates CHD4 recruitment to postnatal-specific regulatory regions

To determine whether NFIA facilitates CHD4 recruitment to postnatal-specific genomic regions, we first performed NFIA CUT&RUN in wild-type neonatal cardiomyocytes. CUT&RUN-qPCR analysis across the regulatory regions previously identified by CHD4 binding (Figures 4C and S4B, C) confirmed robust NFIA occupancy at all tested regions of representative target genes, including *Slc27a1*, *Lgr6*, and *Dgat2* (Figure 6E). We next investigated whether CHD4 occupancy at these regions depends on NFIA. For this purpose, we isolated neonatal cardiomyocytes from *Nfia^flox/flox^* mice at P0 and treated them with *Cre*-expressing adenovirus or control GFP-expressing adenovirus (Figure 6F), achieving efficient NFIA deletion (Figure 6G). Subsequent CHD4 CUT&RUN analysis revealed that NFIA deletion did not substantially affect CHD4 occupancy at stage-independent binding regions, such as Region 2 of *Slc27a1* and *Dgat2*, which are occupied by CHD4 in both embryonic and postnatal hearts (Figure 6H, K, and 4C). In contrast, CHD4 occupancy was reduced at postnatal-specific binding regions, including Regions 1 and 3 of *Slc27a1* and Region 1 of *Dgat2* (Figure 6I, J, L). Together, these findings demonstrate that NFIA facilitates CHD4 occupancy at selected postnatal-specific genomic regions and contributes to the expression of a subset of genes involved in cardiac fatty acid metabolism. These results identify NFIA as a developmentally regulated CHD4-interacting factor that helps establish the postnatal CHD4-dependent transcriptional program in cardiomyocytes.

## DISCUSSION

Although CHD4 is recognized as a transcriptional repressor during cardiac development and homeostasis, roles for gene program activation are emerging. Here, we demonstrate CHD4 is required for promoting the postnatal fatty acid metabolic gene program and identify NFIA as a developmentally regulated factor that facilitates this activity by regulating CHD4 genomic occupancy. First, our comparative transcriptomic analyses demonstrate that CHD4 has a prominent role in activating fatty acid metabolic genes in postnatal cardiomyocytes (Figures 2 and 3). Second, we identify developmental stage-dependent differences in CHD4 chromatin occupancy, including increased occupancy at binding regions associated with metabolic genes in postnatal cardiomyocytes (Figures 4 and S4). Third, we establish a physical association between CHD4 and NFIA in postnatal cardiomyocytes and demonstrate that NFIA facilitates CHD4 occupancy at selected postnatal-specific regulatory regions (Figures 5 and 6). Together, these findings reveal an NFIA-CHD4 regulatory axis that contributes to the transcriptional control of fatty acid metabolism during postnatal cardiac maturation.

Consistent with previous findings by Gomez-del Arco et al., postnatal cardiomyocyte-specific deletion of *Chd4* resulted in severe cardiac dysfunction, myofibrillar disassembly, abnormal mitochondrial cristae morphology, and premature mortality^17^. Our findings extend this work by identifying a previously unrecognized gene-activating function of CHD4 in postnatal cardiac metabolic maturation. Although CHD4 regulates cardiac development and function at multiple developmental stages^15–22^, comparative transcriptomic analyses reveal that its contribution to fatty acid metabolic gene expression is specific to the postnatal heart (Figures 2 and 3). The activation of the metabolic gene program in postnatal cardiomyocytes was associated with increased CHD4 genomic occupancy (Figures 4 and S4), suggesting CHD4 functions as a transcriptional activator. Similarly, a set of genes involved in fatty acid uptake and intracellular lipid handling, including *Cd36*, *Slc27a1*, *Dgat2*, *Fabp3*, and *Lpl*, as well as mitochondrial genes such as *Ndufa8*, *Mtfp1*, and *Mtg1*, were downregulated in postnatal *Chd4* knockouts, further demonstrating CHD4 functions as a transcriptional activator, and linking CHD4 transcriptional regulation to defects in lipid-handling and mitochondrial function observed in *Chd4* knockouts (Figures 2 and S2). In addition, the redistribution of lipids from intracellular droplets to the cardiac interstitium is consistent with impaired lipid handling by mutant cardiomyocytes (Figure 2G, H). Collectively, these findings expand the established functions of CHD4 in the heart by revealing a new transcriptional-activating role to control postnatal fatty acid metabolic gene program.

Our study also reveals NFIA as a dynamic transcription factor with increased postnatal expression, that interacts with CHD4 to activate metabolic programs. NFIA protein expression was highest in cardiomyocytes during the early postnatal period and markedly reduced in adult hearts (Figure 5A), suggesting it functions in establishing postnatal metabolic maturation, but might be dispensable at later stages. Reciprocal immunoprecipitation, mass spectrometry, and proximity ligation demonstrate NFIA and CHD4 directly interact within neonatal cardiomyocyte nuclei. Mechanistically, CUT&RUN analyses revealed that NFIA promotes CHD4 occupancy at specific regions associated with metabolic genes, but not other regions. Although *Nfia*-deficient hearts exhibit a milder phenotype than *Chd4*-deficient hearts, genes downregulated in both models included a subset of metabolic genes. Differences in phenotypic severity of *Nfia* inactivation vs *Chd4* inactivation likely reflect additional non-metabolic gene regulatory functions of CHD4, for example, in repressing expression of the non-cardiomyocyte myofibrillar gene program. Together, these findings demonstrate NFIA facilitates CHD4 recruitment to a series of postnatal-specific regulatory regions to activate metabolic genes during postnatal cardiomyocyte maturation.

Mechanisms underlying CHD4 recruitment might involve additional factors. Our data revealed 17 genes commonly reduced by inactivating either factor, while canonical NFI motifs were not significantly enriched in the CHD4-bound regions associated with the 17 genes, but the PPARγ motif was significantly enriched (Figure S6K). This observation is similar to previously described interactions between NFIA and PPARγ at brown-fat-specific regulatory regions that have been shown to facilitate transcriptional activation^28^. PPARγ also plays an important role in myocardial lipid metabolism, as cardiomyocyte-specific PPARγ overexpression increases fatty acid oxidation gene expression and triglyceride uptake but paradoxically leads to myocardial lipid accumulation, mitochondrial structural abnormalities, and dilated cardiomyopathy^73^. These findings highlight the importance of tightly coordinated PPARγ-dependent transcriptional regulation in maintaining myocardial metabolic homeostasis. Given the enrichment of PPARγ motifs in CHD4-bound regions associated with commonly regulated genes, we speculate that PPARγ may participate in the NFIA–CHD4 regulatory network to coordinate postnatal metabolic maturation in cardiomyocytes. However, whether PPARγ occupies these genomic regions and functionally cooperates with NFIA and CHD4 to regulate metabolic gene expression in cardiomyocytes remains to be determined.

Since perinatal *Chd4* knockouts results prevents metabolic maturation, this model may provide a useful experimental system for linking defects in metabolic maturation to early-onset cardiovascular disease. Future work will determine whether similar metabolic phenotypes are associated with specific forms of pediatric cardiac disease and whether targeting these pathways can improve cardiac function.

Our study has several limitations. First, although we observed reduced expression of genes involved in fatty acid uptake and metabolism, together with altered myocardial lipid distribution, we were unable to directly quantify fatty acid uptake kinetics in primary neonatal cardiomyocytes using commercially available fluorescent fatty acid uptake assays due to insufficient signal sensitivity. Future studies using radiolabeled fatty acid tracers or mass spectrometry-based metabolic flux analysis are required to quantify fatty acid uptake and utilization more precisely. These approaches will also help distinguish the relative contributions of impaired fatty acid uptake, intracellular lipid handling, and mitochondrial utilization to the metabolic phenotype of *Chd4*-deficient hearts.

Second, Oroboros high-resolution respirometry revealed an apparent discrepancy between the suppression of the FAO gene program and mitochondrial respiratory function. Although absolute FAO-supported oxygen consumption was not significantly altered in *Chd4-cmKO* hearts, maximal electron transfer system capacity and Complex IV-supported respiration were reduced. Furthermore, increased flux control ratios for FAO and several substrate-supported respiratory states likely reflected the reduction in maximal respiratory capacity rather than enhanced absolute FAO capacity. These findings suggest that, despite transcriptional suppression of FAO-related genes, mitochondrial capacity for substrate-supported respiration may be partially preserved, whereas the broader mitochondrial respiratory machinery is compromised. Further studies directly measuring fatty acid oxidation flux and mitochondrial ATP production will be necessary to clarify the functional consequences of CHD4 deficiency on myocardial fatty acid utilization and energy metabolism. It remains to be determined whether restoring the expression of key lipid transporters or metabolic genes can rescue *Chd4* inactivation.

Third, cardiomyocyte-specific deletion of *Nfia* resulted in a considerably milder cardiac phenotype than deletion of *Chd4*. Although *Nfia-cmKO* mice exhibited altered cardiac function and downregulation of metabolic genes, they did not develop overt heart failure or premature mortality. A likely explanation is functional compensation by other NFIs, including NFIB, NFIC, and NFIX, which share conserved DNA-binding domains and may recognize overlapping DNA sequences^70,74^. However, their expression in cardiomyocytes and the ability to functionally compensate in *Nfia*-deficient cardiomyocytes remains to be tested. The milder phenotype of *Nfia-cmKO* mice likely also reflects the broader regulatory functions of CHD4, which interacts with multiple transcription factors and chromatin-associated proteins to regulate diverse gene programs.

In conclusion, our findings identify a developmentally regulated NFIA-CHD4 interaction that facilitates CHD4 recruitment to postnatal-specific genomic regions and promotes the activation of a subset of genes involved in fatty acid metabolism. This work expands the understanding of CHD4 beyond its established repressive functions and identifies NFIA as a stage-specific regulator that contributes to CHD4-mediated gene activation during postnatal cardiomyocyte maturation.

## AUTHOR CONTRIBUTIONS

M.C., Y.Y.S., M.E.S., and W.S. conceived the project. M.C., Y.Y.S., and Y-W.C. designed and performed most of the experiments. L.M. performed the bioinformatic analyses for RNA-seq, CHD4 genomic binding shift, and DNA motif mapping. K.P. performed the Oroboros assay and data analysis. The manuscript was written by M.C., Y.Y.S., M.E.S., and W.S., and all authors provided suggestions for improvement.

## ACKNOWLEDGEMENTS

We sincerely thank Dr. Frank Conlon (University of North Carolina at Chapel Hill) for sharing the *Chd4 flox* allele (originally generated by Dr. Katia Georgopoulos’s laboratory at Harvard Medical School), and we greatly thank Dr. Benjamin Deneen (Baylor College of Medicine) for sharing the *Nfia flox* allele. Mass spectrometry was performed using instrumentation in the CardiOmics Program supported by the University of Nebraska Medical Center (UNMC) Center for Heart and Vascular Research. We appreciate the assistance of Dr. Bryan Hackfort at the UNMC Animal Cardiovascular Physiology Core with mouse echocardiographic measurement. We acknowledge use of the UNMC Animal Behavior Core Facility, RRID:SCR_018830, P20GM130447, with additional appreciation to Dr. Mystera Samuelson and Jessica Post of the UNMC Animal Behavior Core for their assistance. We recognize the use of the Electron Microscopy Core Facility (EMCF; RRID:SCR_017864), which receives support from the UNMC Department of Pathology, Microbiology, and Immunology. The graphical schematics in Figures 1A and 6F were generated using Biorender.

## Nonstandard Abbreviations and Acronyms

AAV: adeno-associated virus
CHD4: chromodomain helicase DNA binding protein 4
ChIP: chromatin immunoprecipitation
cTnT: cardiac troponin T
CUT&RUN: cleavage under targets and release using nuclease
EF: ejection fraction
FS: fractional shortening
GSEA: gene set enrichment analysis
IP/MS: immuno-purification/mass spectrometry
NFIA: nuclear factor I A
NuRD: nucleosome remodeling and deacetylation
PPARγ: peroxisome proliferator-activated receptor gamma
TMY: tropomyosin

---

**Figure S1.**
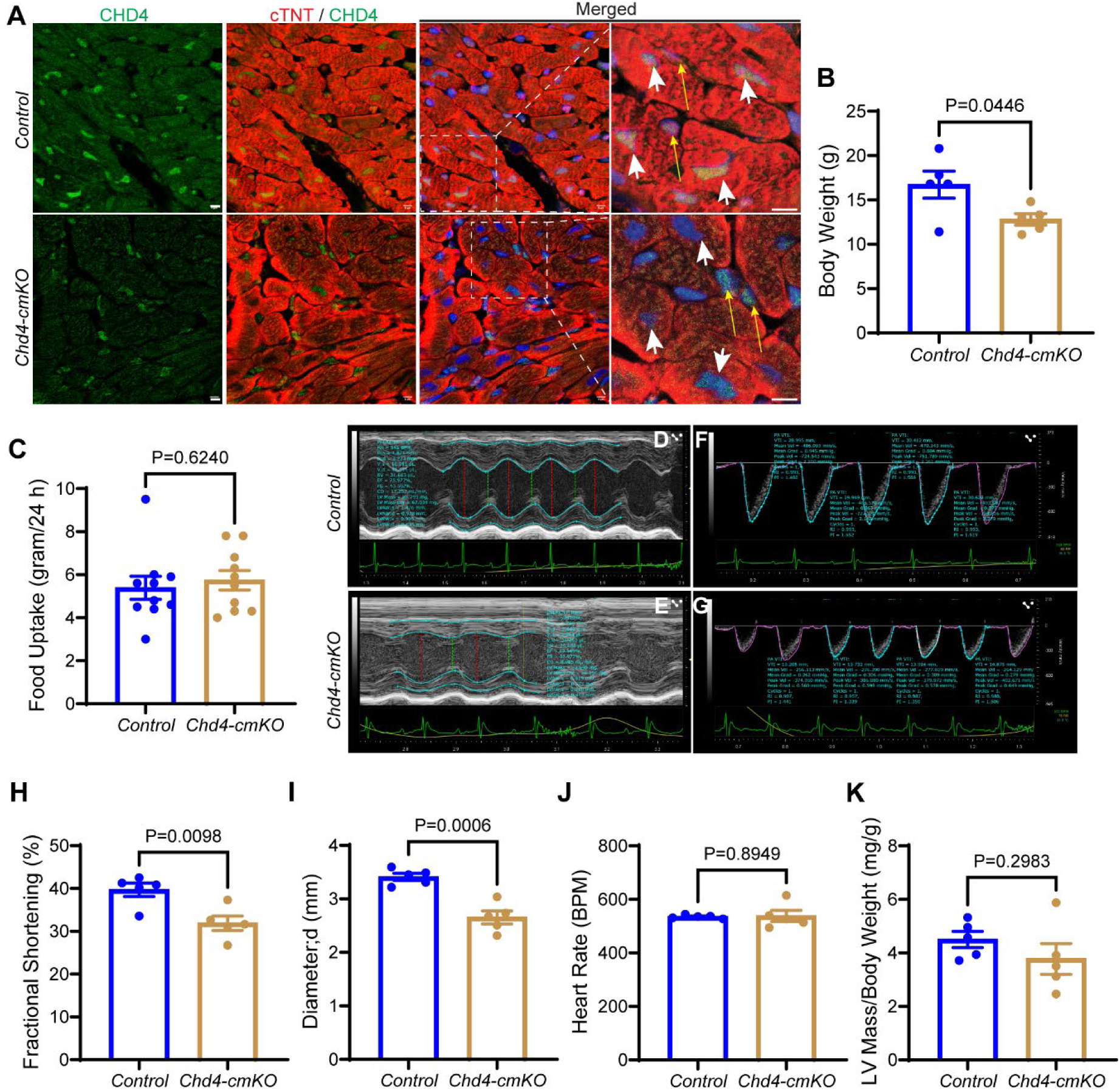
*Chd4* deficiency in postnatal cardiomyocytes results in cardiac dysfunction. (A) Representative images of *Chd4-cmKO* and *Control* heart sections stained with anti-CHD4 (green), anti-cTNT (red), and DAPI (blue). White arrows: cardiomyocyte nuclei. Yellow arrows: non-cardiomyocyte arrows. Scale bars: 5 μm. (B) Body weight of the *Control* and *Chd4-cmKO* mice at four weeks. P value: unpaired Student’s t test. (C) Food consumption of the *Control* and *Chd4-cmKO* mice in 24 hours during the Scurry Activity Wheel assay. P value: unpaired Student’s t test. (D, E) Representative M-mode echocardiograms of left ventricular (LV) systolic function and chamber dimensions in *Control* and *Chd4-cmKO* mice. Red dashed lines indicate left ventricular internal diameter at diastole (LVIDd), green dashed lines indicate left ventricular internal diameter at systole (LVIDs), and bottom green traces show simultaneous electrocardiogram (ECG) recordings. (F, G) Representative pulsed-wave (PW) Doppler spectral tracings of pulmonary artery (PA) outflow in *Control* and *Chd4-cmKO* mice. (H-K) Quantifications of cardiac function parameters. P values: unpaired Student’s t test.

**Figure S2.**
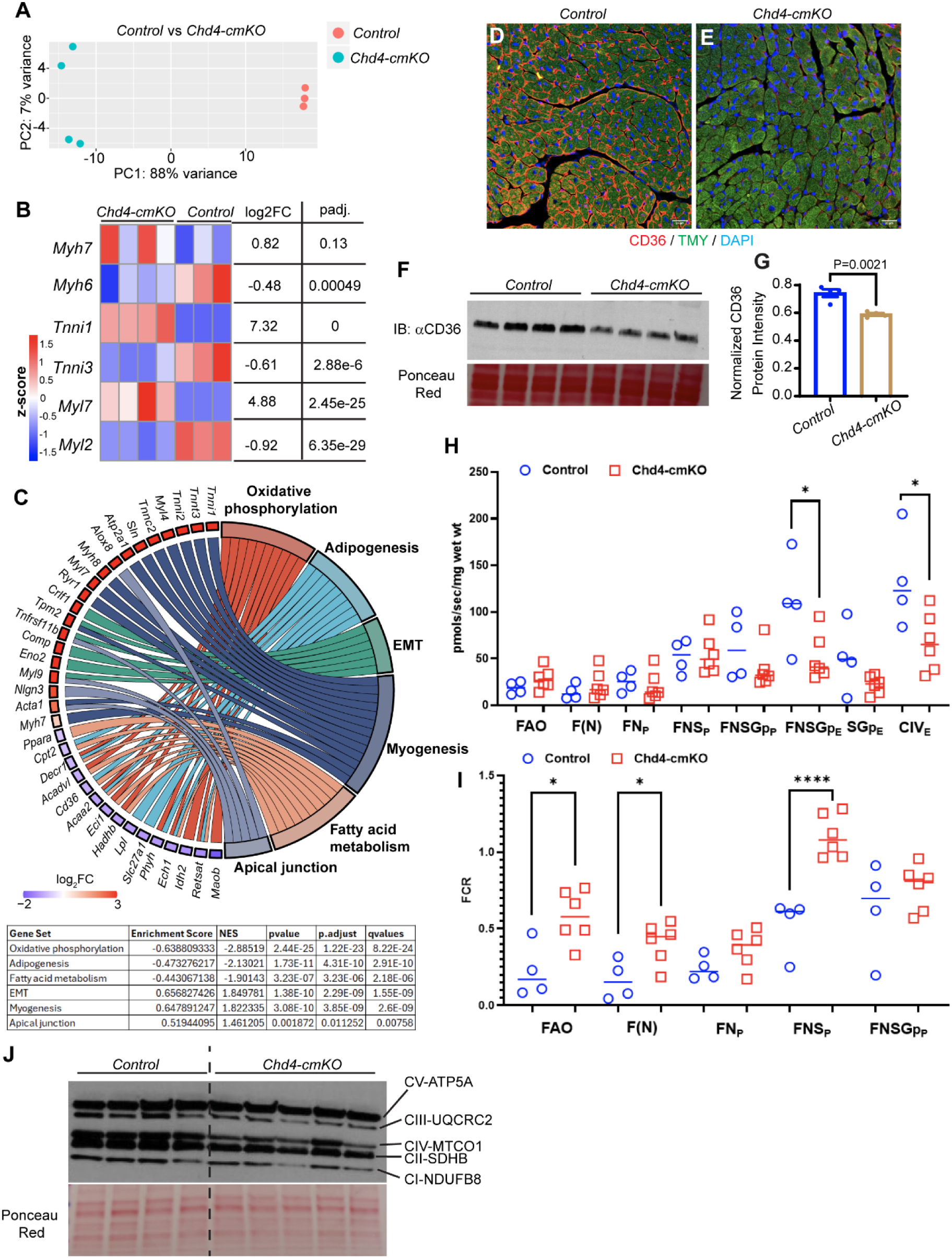
Dysregulated transcriptome in the *Chd4-cmKO* mouse hearts. (A) Principal component analysis (PCA) plot of gene expression data from RNA-seq of biological replicates for each genotype. (B) Heatmap of selected genes from the transcriptomes. Color bar: z-score. (C) Circular plot of the overrepresented Biological Process GO terms in the *Chd4-cmKO* hearts, simultaneously presenting a detailed view of the relationships between expression changes (left semicircle perimeter) and enriched biological processes (right semicircle perimeter). Color bar: Log_2_(Fold Change) of gene expression. (D, E) Representative immunohistochemistry-stained images of the *Control* and *Chd4-cmKO* heart sections. Red: CD36 (arrows); green: Tropomyosin (TMY), marker of cardiomyocytes; blue: DAPI. Scale bars: 20 μm. (F) Western blot of the whole heart lysate followed by probing anti-CD36 antibody. Before antibody probing, the blot was stained with Ponceau S Red solution to present the protein loading amount in each lane. (G) Quantification of the CD36 protein band intensity in (F), normalized to Ponceau S Red loading control. P value: unpaired Student’s t test. (H) High-resolution respirometry analysis of mitochondrial respiration in *Control* and *Chd4-cmKO* hearts using the Oroboros system. Oxygen consumption rates (OCR) were normalized to wet heart tissue weight (pmol O₂/sec/mg wet wt) and measured under different substrate and inhibitor conditions to assess fatty acid oxidation (FAO), substrate-supported oxidative phosphorylation, maximal electron transfer system (ETS) capacity, and Complex IV (CIV)-supported respiration. *P < 0.05, unpaired Student’s t test. (I) Flux control ratios (FCRs) of mitochondrial respiration in *Control* and *Chd4-cmKO* hearts, calculated by normalizing oxygen consumption rates under individual substrate and inhibitor conditions to FCCP-induced maximal uncoupled respiration. Blue circles, control; red squares, Chd4-cmKO. Horizontal lines indicate group means. *P < 0.05; ****P < 0.0001, unpaired Student’s t test. (J) Representative Western blot analysis of mitochondrial oxidative phosphorylation (OXPHOS) complex subunits in *Control* and *Chd4-cmKO* hearts. Representative subunits of Complex I (NDUFB8), Complex II (SDHB), Complex III (UQCRC2), Complex IV (MTCO1), and Complex V (ATP5A) were examined using an OXPHOS antibody cocktail. The blot was stained with Ponceau S Red solution to present the protein loading amount in each lane.

**Figure S3.**
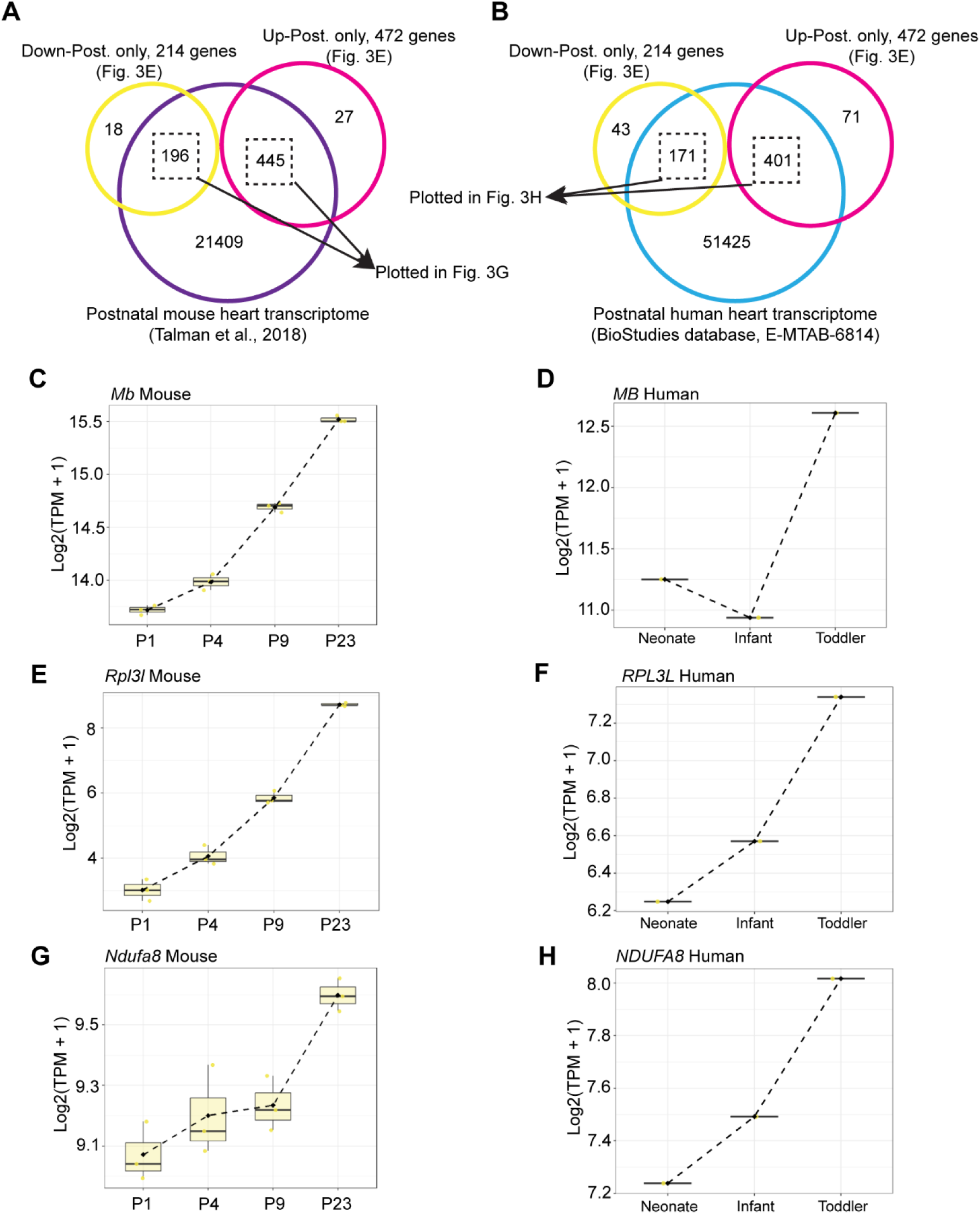
Mouse and human postnatal heart transcriptomes. (A, B) Genes only upregulated in postnatal *Chd4-cmKO* mouse hearts (“Up-Post. only”, N=472 genes) and genes only downregulated in postnatal *Chd4-cmKO* mouse hearts (“Down-Post. only”, N=214 genes) (from Fig. 3E) were overlapped to a wildtype postnatal mouse heart transcriptomic dataset across four stages: postnatal day 1 (P1), P4, P9, and P23 (Talman et al., 2018) (A) and to a human heart transcriptomic dataset of three postnatal stages: neonate, infant, and toddler (BioStudies database (http://www.ebi.ac.uk/biostudies) under accession number E-MTAB-6814) (B). 445 of the 472 “Up-Post. only” genes and 196 of the 214 “Down-Post. only” genes were found in the Talman mouse dataset (A) and 401 of the 472 “Up-Post. only” genes and 171 of the 214 “Down-Post. only” genes were found in the BioStudies human dataset (B). Overlapped genes were plotted in Fig. 3F and 3G as indicated. (C-H) Expression dynamics of representative genes in the mouse and human postnatal heart datasets.

**Figure S4.**
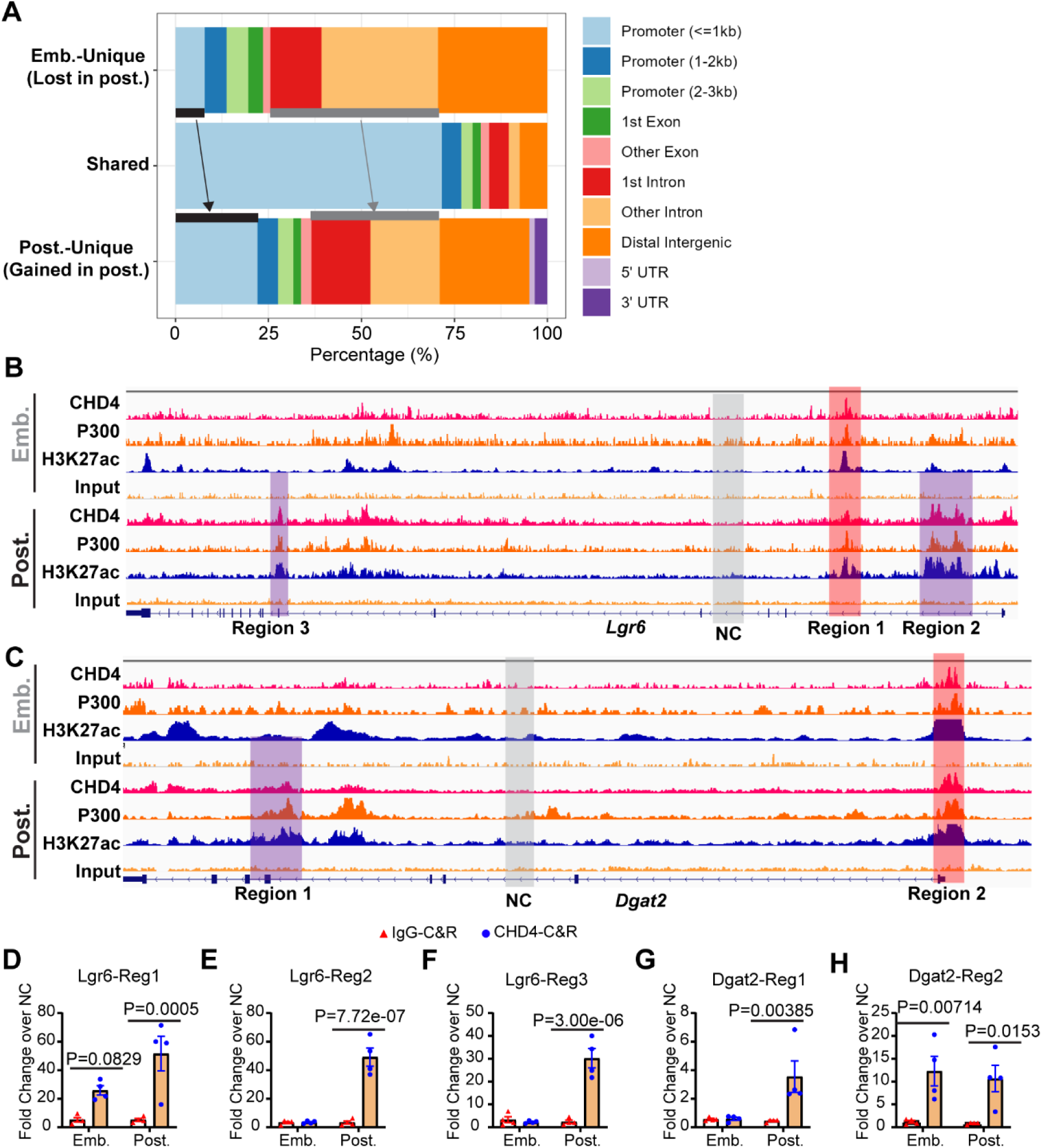
CHD4-bound regions emerging in the postnatal heart. (A) Genomic feature annotation of dynamic CHD4 binding sites. Genomic distribution of embryonic-unique (lost), shared (conserved), and postnatal-unique (gained) CHD4 binding peaks across functional annotation categories (promoters, exons, introns, and distal intergenic regions). Conserved binding sites (Shared) are predominantly enriched at promoters (<= 1kb), whereas stage-specific dynamic peaks (Emb.-Unique and Post.-Unique) show a broader distribution across distal intergenic regions and intronic elements, reflecting stage-specific shifts in regulatory genomic architecture. (B, C) Representative Integrative Genomic Viewer (IGV) browser tracks of the CHD4-bound regions on the *Lgr6* and *Dgat2* genes. Regions highlighted in red are bound by CHD4 in both embryonic and postnatal hearts, whereas Regions highlighted in purple represent postnatal binding sites. A Region with few reads in both embryonic and postnatal CHD4-ChIP-seq tracks were selected as the negative control (NC) region (highlighted in grey) used for CHD4-CUT&RUN (Cleavage Under Targets & Release Using Nuclease) enrichment calculation. ChIP-seq tracks of the active histone mark H3K27ac and the histone acetyltransferase P300 in embryonic and postnatal mouse hearts were used to indicate the active transcriptional feature of highlighted regions. ChIP-seq data were retrieved from literature. CHD4-ChIP-seq in the embryonic mouse heart (E10.5): Wilczewski et al, 2018 (GSE109012). CHD4-ChIP-seq in the postnatal mouse heart (atria) (P24): Sweat et al., 2025 (GSE284095). P300-ChIP-seq in the embryonic (E12.5) and postnatal (P28) mouse heart: Zhou et al., 2023 (GSE195901). H3K27ac-ChIP-seq in the embryonic mouse heart (E10.5): Gu et al., 2022 (GSE213572). H3K27ac-ChIP-seq in the postnatal mouse heart (atria) (P28): Sweat et al., 2023 (GSE222970). (D-H) Cardiomyocytes were isolated from wildtype C57Bl/6 mouse hearts at E10.5 or P1-P2 and cultured for 72 hours. CHD4-and IgG-CUT&RUN were performed using the isolated cardiomyocytes followed by library preparation and region-targeted quantitative PCR (qPCR). Genomic DNA purified from isolated embryonic or neonatal cardiomyocytes, without antibody targeting, was used as an input DNA to normalize the enrichment signal and the selected NC Region (indicated in Fig. S4B, C) was used to calculate the enrichment fold of targeted Regions. P values: two-way ANOVA.

**Figure S5.**
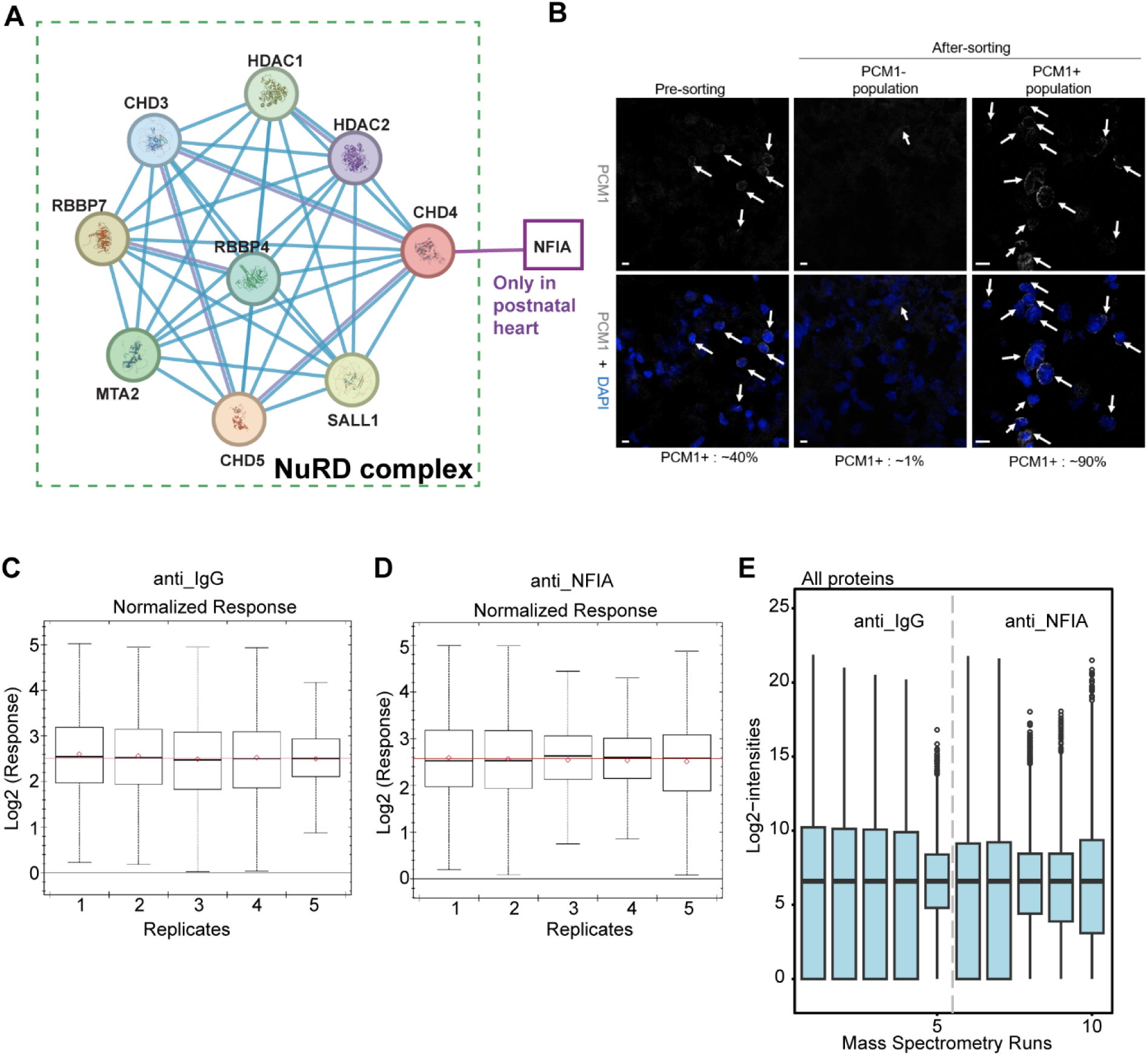
Quality control and normalization of mass spectrometry interactome data for anti-IgG and anti-NFIA IP/MS. (A) Functional protein association networks of CHD4 cardiac interactome in the postnatal mouse hearts. The postnatal CHD4 cardiac interactome is from Shi et al., Development, 2024. (B) Representative immunofluorescent images of neonatal cardiac nuclei stained with anti-PCM1 (pericentriolar material 1 protein) and DAPI. Nuclei were isolated from wildtype C57Bl/6 mouse hearts at P1-P2. The isolated nuclei were then incubated with anti-PCM1 antibody followed by Magnetic-activated cell sorting (MACS). Pre-sorting and after-sorting nuclei were then stained with anti-PCM1 antibody for fluorescent imaging. The PCM1^+^ nuclei were used for the NFIA (and IgG)-IP/MS assays. (C, D) Box plots showing normalized Log2 response distributions across five technical replicates for control IgG (C) and anti-NFIA (D) samples, demonstrating consistent median values and overall signal distribution across runs. (E) Box and whisker plots of Log2 protein intensities across all ten mass spectrometry runs (five anti-IgG and five anti-NFIA runs), illustrating uniform global intensity distributions and stable technical reproducibility across the experimental dataset.

**Figure S6.**
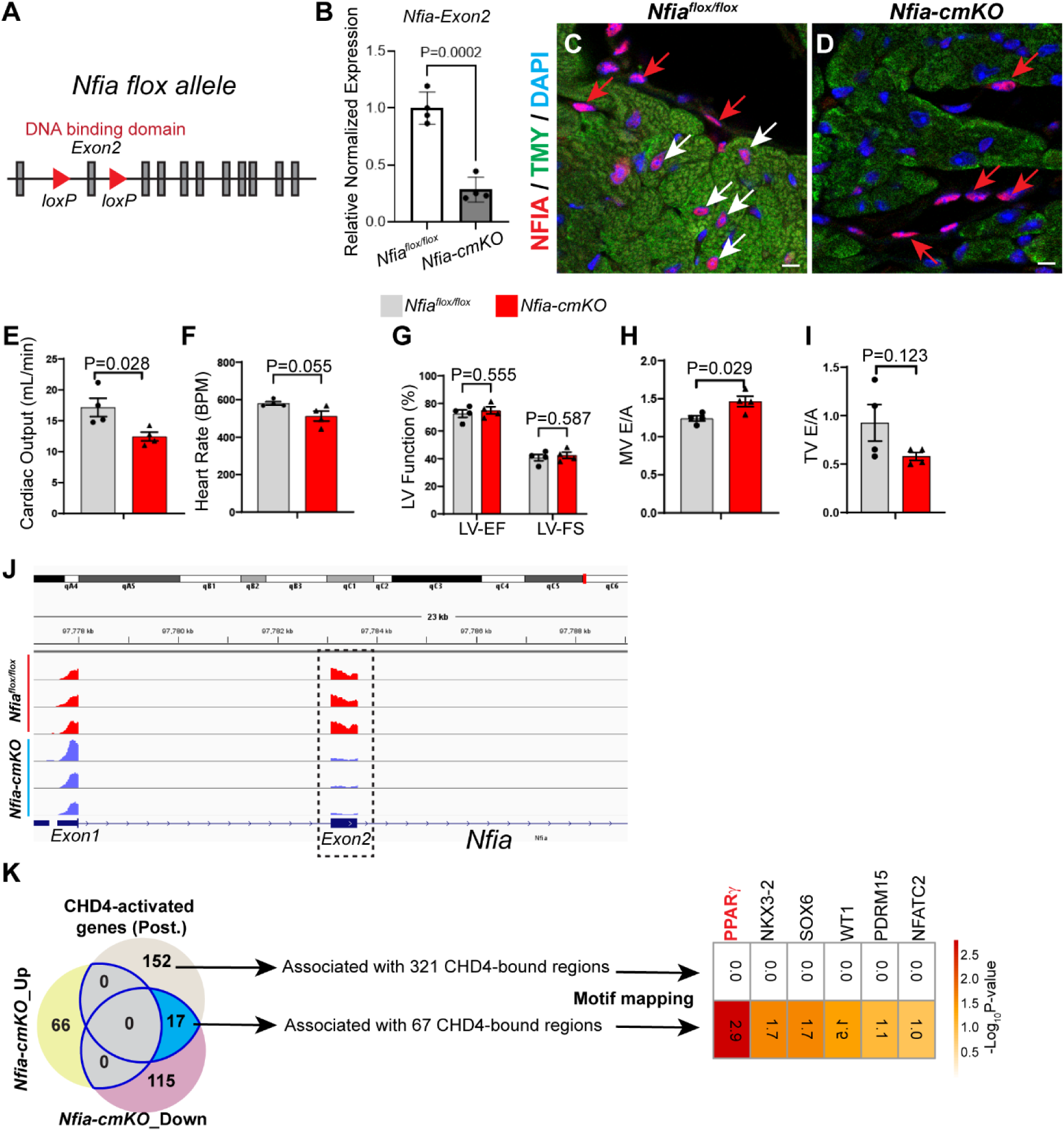
*Nfia* deficiency in cardiomyocytes. (A) Schematic representation of *Nfia^flox^* allele, with loxP sites flanking the second exon containing the DNA binding domain. (B) *Nfia* exon 2-targeted RT-qPCR with the *Nfia^flox/flox^* control and *Nfia-cmKO* mouse hearts. The *Rpl32* gene was used as internal control, and gene expression in the control hearts was normalized as 1.0. P values: unpaired Student’s t test. (C, D) Representative immunohistochemistry-stained images of the *Nfia^flox/flox^*control and *Nfia-cmKO* heart sections. Red: NFIA (arrows); green: Tropomyosin (TMY), marker of cardiomyocytes; blue: DAPI. White arrows: NFIA^+^ cardiomyocytes. Red arrows: NFIA^+^ noncardiomyocytes. Scale bars: 10 μm. (E-I) Quantifications of cardiac function parameters of the *Nfia^flox/flox^* control and *Nfia-cmKO* mice at 4 weeks old. P values: unpaired Student’s t test. (J) Representative Integrative Genomic Viewer (IGV) browser tracks of *Nfia* gene in the RNA-seq of the *Nfia^flox/flox^* control and *Nfia-cmKO* mouse hearts. Decreased reads of exon 2 is highlighted. (K) Genomic intersection and transcription factor motif enrichment at CHD4–NFIA co-regulated distal enhancers. Venn diagram illustrating the overlap between postnatal CHD4-activated genes and genes differentially expressed in *Nfia-cmKO* hearts. The intersection highlights 17 core co-activated target genes down-regulated in *Nfia-cmKO* hearts that are directly regulated by CHD4 (blue outline). Genomic regions bound by CHD4 within these gene sets (321 regions associated with the 152 CHD4-activated non-overlapping genes and 67 regions associated with the 17 co-regulated genes) were subjected to motif enrichment analysis. Heatmap displays-Log_10_(P value) enrichment of transcription factor motifs mapped using the monaLisa R package against the JASPAR database, revealing selective and statistically significant enrichment of the PPARγ binding motif (as well as NKX3-2, SOX6, WT1, PRDM15, and NFATC2) exclusively at the 67 CHD4-bound regions corresponding to the 17 co-regulated gene cassette.

